# Longitudinal deep phenotyping in a mouse model of West syndrome reveals temporal dynamics of synapse remodeling, gliosis, and proteomic and lipidomic changes during seizure evolution

**DOI:** 10.1101/2025.05.06.652352

**Authors:** Ruiying Ma, Muwon Kang, Gyu Hyun Kim, Hyojin Kang, Sunho Lee, Yeji Yang, Hae Ji Lee, Seungji Choi, Seungsoo Kim, Seoyeong Kim, Yukyung Jun, Hyewon Kim, Yinhua Zhang, U Suk Kim, Hyae Rim Kang, Yoonhee Kim, Yulim Lee, Woosuk Chung, Eunha Kim, Minji Jeon, Geum-Sook Hwang, Jungmin Choi, Youngae Jung, Jin Young Kim, Eunjoon Kim, Kea Joo Lee, Kihoon Han

## Abstract

Neurodevelopmental disorders can have long-lasting effects, causing not only early pediatric symptoms but also a range of neurological issues throughout adulthood. West syndrome is a severe neurodevelopmental disorder marked by infantile spasms, an early symptom that typically subsides with age. However, many patients progress to other seizure forms, known as seizure evolution, which is closely linked to poor long-term outcomes. Despite its clinical significance, the neurobiological mechanisms behind seizure evolution in West syndrome remain poorly understood. Recent genetic studies have consistently identified the *CYFIP2* p.Arg87Cys variant in West syndrome patients, and the *Cyfip2^+/R87C^* mouse model carrying this mutation has been shown to recapitulate key symptoms of the disorder, including infantile spasms. In this study, we aimed to gain deeper insight into seizure evolution by conducting longitudinal deep phenotyping of the *Cyfip2^+/R87C^* mouse model from the neonatal stage to seven months of age. We tracked seizure activity through behavioral and EEG recordings and employed multi-omic analyses, including tissue and single-cell level transcriptomics, ultrastructural analysis, proteomics, and lipidomics, to capture a comprehensive view of molecular and cellular changes. Our results showed that after an initial period of neonatal spasms, *Cyfip2^+/R87C^* mice entered a seizure-free phase, followed by spontaneous recurrent seizures in adulthood, ultimately leading to premature death. This progression was associated with synaptic remodeling, sequential activation of different glial cell types, lipid droplet accumulation in astrocytes, and significant proteomic and lipidomic changes in the brain. These findings suggest that seizure evolution in West syndrome involves complex, time-dependent interactions between neurons and glial cells, along with alterations in lipid metabolism. Our study highlights the potential of longitudinal multi-omic approaches to uncover underlying mechanisms of seizure evolution and suggests that targeting these changes could offer novel therapeutic strategies. Additionally, the dataset generated here may provide valuable insights for other epilepsy and neurodevelopmental disorder models.

## Introduction

Neurodevelopmental disorders and pediatric neurological conditions often lead to premature death and long-term disabilities, posing a significant burden on society [1]. One such example is West syndrome, also known as infantile spasms [2]. It is a rare but severe neurodevelopmental disorder, occurring in 2 to 3 out of every 10,000 live births [3]. It is characterized by spasms, hypsarrhythmia on electroencephalogram, and developmental regression, typically presenting within the first year of life. While the prognosis varies depending on factors such as etiology and degree of developmental regression, the mortality rate of West syndrome can reach up to 40% depending on the length of follow-up [4]. Moreover, even after the initial spasms subside, up to 60% of patients go on to develop other seizure types, including Lennox-Gastaut syndrome, which are associated with poor long-term outcomes [5]. Notably, sudden unexpected death in epilepsy (SUDEP) remains a significant cause of mortality in these patients [6]. Despite the clinical significance, the detailed neurobiological mechanisms underlying seizure evolution in West syndrome remain poorly understood. Understanding these mechanisms could uncover potential targets to prevent seizure progression.

To better comprehend these mechanisms, particularly at the molecular and cellular levels, it is crucial to have an animal model that reflects the same underlying etiology [7, 8], such as a genetic mutation identified in patients, and that exhibits both the hallmark symptoms of West syndrome and the progression of seizure evolution. Furthermore, conducting longitudinal deep phenotyping of the animal model, from the stages of spasms to the seizure-free latent period and onward to the onset and progression of different seizure types, is key to gaining a comprehensive understanding of the seizure evolution process. To the best of our knowledge, however, such an extensive study of West syndrome models has not yet been reported.

Cytoplasmic FMR1-interacting protein 2 (CYFIP2) is an evolutionarily conserved protein that is highly expressed in the brain [9]. At the molecular level, CYFIP2 is a key component of the Wiskott–Aldrich syndrome protein family verprolin-homologous protein (WAVE) regulatory complex, which regulates actin polymerization and branching [10]. Moreover, CYFIP2 interacts with various RNA-binding proteins and components of membraneless organelles, such as stress granules, implicating its involvement in mRNA processing and translation [11, 12]. Consistently, alterations in CYFIP2 expression lead to neuronal abnormalities, as demonstrated in cultured neurons and mouse models [13–16].

Recent whole exome and genome sequencing studies have identified *de novo CYFIP2* variants in pediatric patients with early-onset epileptic encephalopathy and developmental regression (developmental and epileptic encephalopathies 65, DEE65) [2, 17–19]. Notably, approximately 40% of these cases involve substitutions at the Arg87 “hotspot” residue of CYFIP2, such as Arg87Cys, Arg87Leu, and Arg87Pro [19]. Patients with these hotspot variants often present with more severe symptoms and are predominantly diagnosed with West syndrome [20, 21]. In addition, we recently developed a mouse model carrying the *Cyfip2* p.Arg87Cys variant (*Cyfip2^+/R87C^* mice), which recapitulates many of the clinical features observed in patients, including neonatal spasms and developmental delays [22]. These findings confirm the causal role of the *CYFIP2* p.Arg87Cys variant in the development of West syndrome and the relevance of the *Cyfip2^+/R87C^* mouse model.

Despite these advancements, systemic clinical data on the long-term prognosis of patients carrying *CYFIP2* p.Arg87 variants remain limited, as these cases have only been reported within the last six to seven years. To address this gap, our study was designed in two key stages. First, by observing survival and seizure onset in *Cyfip2^+/R87C^* mice over an extended period, we aimed to determine whether and when seizure evolution occurs in this model. Our findings reveal that *Cyfip2^+/R87C^* mice, following a seizure-free period after neonatal spasms, develop spontaneous recurrent seizures in adulthood, ultimately leading to premature death. Second, based on the temporal data obtained from the seizure evolution process, we conducted multi-omic analyses, including tissue and single-cell level transcriptomics, ultrastructural analysis, proteomics, lipidomics, and their subsequent validation. This comprehensive approach allowed us to gain an in-depth understanding of the molecular and cellular changes involved. We present evidence that seizure evolution in these mice is accompanied by dynamic changes in neuronal synapses, sequential activation of various glial cell types, and significant shifts in the brain proteome and lipidome.

Taken together, these results suggest that multiple cell types and signaling pathways contribute to seizure evolution in West syndrome, emphasizing the potential for developing targeted therapeutic strategies. Moreover, our study represents the first longitudinal deep phenotyping of seizure evolution in a West syndrome model, and the multi-omic datasets generated provide valuable resources for future research. These data provide a foundation for exploring seizure progression mechanisms and can be applied to other models with different etiologies, epilepsy types, or neurodevelopmental disorders, potentially uncovering fundamental mechanisms relevant to a broader range of conditions.

## Results

### Spontaneous recurrent seizures and lethality in adult *Cyfip2^+/R87C^* mice

In our previous study [22], we observed that *Cyfip2^+/R87C^* mice exhibited spontaneous spasms during the neonatal stage (postnatal day 8, PD 8). However, long-term video-EEG recordings from the same study revealed no significant seizures in these mice during the juvenile stage (PD 30), indicating a seizure-free period following neonatal spasms. Unexpectedly, when maintaining *Cyfip2^+/R87C^* mice in their home cages for longer periods, we found that lethality began around postnatal week 14 (PWK 14), with approximately 60% of the mice dying by PWK 30 (**Fig 1A**). In contrast, no significant lethality was observed in *Cyfip2^+/-^* mice, a model for *CYFIP2* haploinsufficiency [15], on either the C57BL/6N or C57BL/6J background. Additionally, *Cyfip2^+/R87C^* mice, but not *Cyfip2^+/-^*, displayed significantly smaller body size and lower body weight at PWK 28 compared to their respective wild-type (WT) littermates (**S1 Fig**).

**Fig 1.**
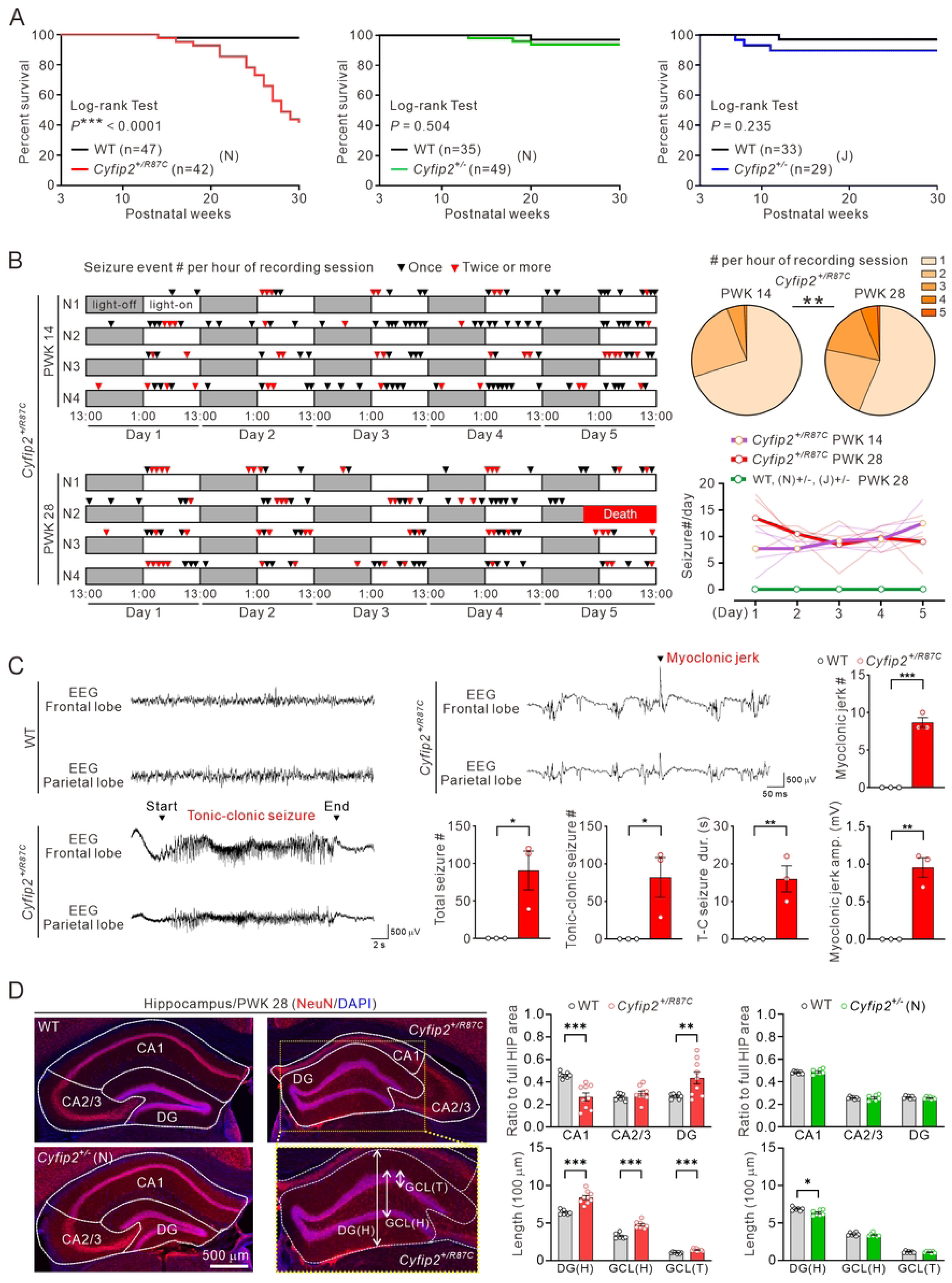
Lethality and spontaneous recurrent seizures in adult *Cyfip2^+/R87C^* mice. (A) Survival curves of *Cyfip2^+/R87C^* mice (on the C57BL/6N (N) background), *Cyfip2^+/-^* mice (on either the C57BL/6N or C57BL/6J (J) backgrounds), and their respective wild-type (WT) littermates, tracked from postnatal weeks 3 to 30 (log-rank test). (B) Behavioral seizure events during long-term (5-day) video recordings of *Cyfip2^+/R87C^* mice at postnatal weeks 14 (PWK 14) and 28 (PWK 28) (each group includes four mice, N1 to N4). Arrowheads indicate one-hour recording sessions with either one (black) or multiple (red) seizure events. *Cyfip2^+/R87C^* mouse N2 of PWK 28 died on day 5 of recording. The pie charts show the distribution of the number of seizure events per one-hour session at PWK 14 and PWK 28 (Fisher’s exact test). The line chart displays the number of seizure events during the 5-day recording for each genotype. Thin lines represent individual mice, while thick lines indicate the group averages. Note that no seizure events were observed in WT or *Cyfip2^+/-^* mice at PWK 28 (n = 4 mice per genotype). (C) Representative traces of EEG recordings from WT and *Cyfip2^+/R87C^* mice at PWK 20. Graphs show quantifications of the total number of seizures, tonic-clonic (T-C) seizures, and myoclonic jerks, as well as the duration (dur.) of T-C seizures and the amplitude (amp.) of myoclonic jerks (n = 3 mice per genotype, unpaired two-tailed Student’s t-test). (D) Representative fluorescence immunohistochemistry images and quantification show a reduction in the CA1 region and dispersion of the dentate gyrus (DG) granule cell layer (GCL) in the hippocampus of *Cyfip2^+/R87C^* mice compared to WT mice at PWK 28 (n = 7 to 9 mice per genotype, two-way ANOVA with Šídák’s multiple comparisons test). *Cyfip2^+/-^* mice displayed a normal hippocampus, except for a reduction in DG height (H) compared to WT mice. T = thickness. \**P* < 0.05; \*\**P* < 0.01; \*\*\**P* < 0.001. Data are represented as mean ± standard error of the mean.

To investigate whether the lethality in *Cyfip2^+/R87C^* mice is associated with their seizures, we conducted long-term video recordings with different cohorts and found that these mice indeed experienced spontaneous recurrent seizures at both PWK 14 and 28 (**Fig 1B and S1 and S2 Videos**). Notably, once they occurred, behavioral seizure events were more clustered at PWK 28 compared to PWK 14, as evidenced by an increased number of seizure events per hour during the recording sessions at PWK 28. In contrast, no such behavioral seizures were observed in WT or *Cyfip2^+/-^* mice at PWK 28. We also attempted simultaneous video-EEG recordings on *Cyfip2^+/R87C^* mice at PWK 20 with different cohorts; however, due to a high mortality rate (∼80%) during or after surgery, likely linked to their poor physical condition, we were unable to analyze a significant number of mice. Nevertheless, in the few surviving *Cyfip2^+/R87C^* mice (n = 3), we observed tonic-clonic seizures and myoclonic jerks that correlated with behavioral seizures, events that were never observed in WT mice (**Fig 1C and S3 and S4 Videos**). Furthermore, consistent with their recurrent seizures, we observed hippocampal pathology in *Cyfip2^+/R87C^* mice at PWK 28, including shrunken CA1 regions and dispersion of dentate gyrus granule cells, hallmarks of epilepsy [23], while these changes were absent in *Cyfip2^+/-^* mice (**Fig 1D**).

These findings suggest that *Cyfip2^+/R87C^* mice undergo seizure evolution, transitioning from neonatal spasms to adult spontaneous recurrent seizures, with the latter being associated with their lethality. Additionally, the absence of this phenotype in *Cyfip2^+/-^* mice indicates that the hotspot p.Arg87 variant likely induces a toxic gain-of-function, rather than merely destabilizing CYFIP2 proteins [22, 24].

### Brain transcriptomic signatures in *Cyfip2^+/R87C^* mice during seizure evolution

Building upon the above findings, we explored longitudinal changes in the brains of *Cyfip2^+/R87C^* mice during seizure evolution. We also analyzed *Cyfip2^+/-^* mice in parallel to validate the toxic gain-of-function effect caused by the p.Arg87Cys variant at the gene expression level. Bulk-tissue transcriptomic analyses were conducted in the frontal cortex and hippocampus of *Cyfip2^+/R87C^* mice, *Cyfip2^+/-^* mice, and their respective WT littermates (all on the C57BL/6N background) at three postnatal stages (PWK 1, 7, and 14), covering the period from neonatal spasms to the onset of lethality and seizures in *Cyfip2^+/R87C^* mice. The two brain regions were selected due to their high CYFIP2 expression, our previous findings which demonstrated morphological and functional changes in neurons within these regions in various *Cyfip2* mouse models [15, 16, 19], and the above finding (**Fig 1D**) showing cytoarchitectural changes in the hippocampus of *Cyfip2^+/R87C^* mice.

In *Cyfip2^+/R87C^* mice, we observed a dramatic increase in differentially expressed genes (DEGs) in both brain regions as the mice aged, particularly in the hippocampus, with 20, 41, and 3,755 DEGs identified at the three stages, respectively (**Fig 2A and S1 Table**). Gene Ontology (GO) analysis of the DEGs at PWK 14, using the Database for Annotation, Visualization, and Integrated Discovery (DAVID), revealed significant terms, notably the downregulation of synapse-related terms (glutamatergic/cholinergic/dopaminergic synapses) and upregulation of extracellular matrix (ECM)-related terms, suggesting substantial synaptic and cytoarchitectural remodeling. In contrast, the number of DEGs in *Cyfip2^+/-^* mice decreased with age, with only 19 DEGs detected in the hippocampus at PWK 14 (**Fig 2B and S2 Table**).

**Fig 2.**
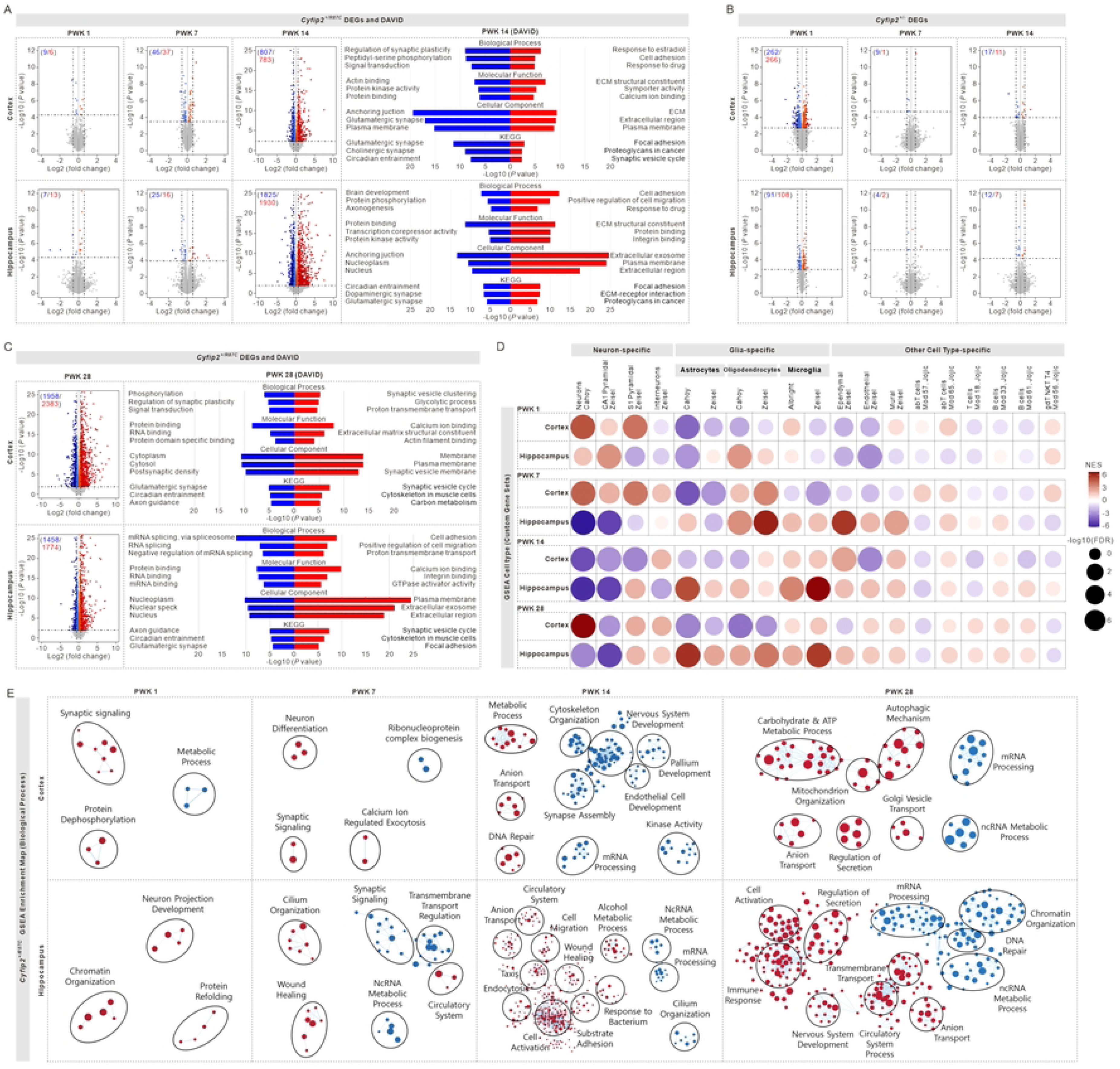
Transcriptomic signatures in the cortex and hippocampus of *Cyfip2^+/R87C^* mice during seizure evolution. (A) Volcano plots display downregulated (blue) and upregulated (red) differentially expressed genes (DEGs) in the cortex and hippocampus of *Cyfip2^+/R87C^* mice compared to WT mice at PWK 1, 7, and 14. Bar graphs present significant terms and their corresponding *P* values from the DAVID (Database for Annotation, Visualization, and Integrated Discovery) Gene Ontology (GO) analysis of the DEGs at PWK 14 (analyses for PWK 1 and 7 were not conducted due to the small number of DEGs). KEGG = Kyoto Encyclopedia of Genes and Genomes. (B) Volcano plots display DEGs in the cortex and hippocampus of *Cyfip2^+/-^* mice at PWK 1, 7, and 14. (C) Volcano plots display DEGs in the cortex and hippocampus of *Cyfip2^+/R87C^* mice at PWK 28. Bar graphs present significant terms and their corresponding *P* values from the DAVID GO analysis of the DEGs. (D) Gene Set Enrichment Analysis (GSEA) of the transcriptome from the cortex and hippocampus of *Cyfip2^+/R87C^* at PWK 1, 7, 14, and 28, focusing on neuron-, glia-, and other cell-type-specific gene sets. NES = normalized enrichment score, FDR = false discovery rate. (E) GSEA of the transcriptome from the cortex and hippocampus of *Cyfip2^+/R87C^* mice, focusing on biological process gene sets, along with clustering of the enriched gene sets using the Cytoscape EnrichmentMap.

We also performed bulk-tissue transcriptomic analysis on the frontal cortex and hippocampus of *Cyfip2^+/R87C^* mice at PWK 28, when their behavioral seizure events were more severe compared to PWK 14. The number of DEGs, particularly in the cortex, was even higher at PWK 28 than at PWK 14 (**Fig 2C and S3 Table**). DAVID GO analysis revealed some overlapping terms between PWK 28 and 14 in both brain regions. However, unlike at PWK 14, RNA regulation-related terms, such as RNA splicing and RNA binding, were significantly downregulated, while synaptic vesicle (i.e., presynapse)-related terms were more prominently upregulated at PWK 28. Terms related to general and postsynaptic functions, such as glutamatergic synapse and postsynaptic density, remained downregulated, which may suggest an imbalance between pre- and postsynaptic compartments, as further validated below.

We also performed Gene Set Enrichment Analysis (GSEA) to identify transcriptomic signatures based on coordinated transcriptional changes across many genes [25]. In testing for cell-type-specific gene sets, we found that the cortex transcriptome of *Cyfip2^+/R87C^* mice shifted from positive to negative enrichment for neuron-specific gene sets between PWK 1 and 14, but became positive again at PWK 28 (**Fig 2D**). Meanwhile, enrichment for glia-specific gene sets in the cortex varied depending on the three glial cell types (i.e., astrocytes, oligodendrocytes, and microglia). In the hippocampus of *Cyfip2^+/R87C^* mice, the age-progressive shift from positive to negative enrichment for neuron-specific gene sets followed a similar pattern to that in the cortex, but occurred earlier and persisted until PWK 28, particularly for the CA1 pyramidal neuron gene set. In contrast, overall enrichment for glia-specific gene sets in the hippocampus shifted from negative to positive with each glial cell type showing a distinct peak of positive enrichment: PWK 7 for oligodendrocytes, PWK 14 for microglia, and PWK 28 for astrocytes. Enrichment for other cell-type-specific gene sets was subtle in both brain regions, except for a notable positive enrichment for ependymal gene sets, particularly in the hippocampus at PWK 7.

GSEA of the *Cyfip2^+/R87C^* transcriptome for biological processes supported the findings from the GO analysis of DEGs and the GSEA for cell-type analyses (**Fig 2E**). Neuron- and synapse-related terms were positively enriched at PWK 1 but became negatively enriched with age. Additionally, RNA regulation-related terms were negatively enriched in both the cortex and hippocampus, particularly at PWK 14 and 28. Meanwhile, terms related to tissue remodeling and immune responses, such as wound healing, cell migration, cell activation, response to bacterium, and immune response, became positively enriched, especially in the hippocampus at PWK 14 and 28.

### Synapse remodeling in *Cyfip2^+/R87C^* mice during seizure evolution

Given the significant age-dependent changes in synapse-related terms observed in the transcriptomic analyses of *Cyfip2^+/R87C^* mice, we sought to validate these findings through additional approaches. qRT-PCR analysis confirmed that the expression levels of genes encoding synaptic proteins, selected from the DEG lists of PWK 14 and 28, notably increased during early postnatal stages but decreased at later stages in the cortex and hippocampus of *Cyfip2^+/R87C^* mice compared to WT mice (**Fig 3A**). In contrast, none of these genes showed significant changes in the cortex and hippocampus of *Cyfip2^+/-^* mice at PWK 28 (data not shown). To further assess synaptic changes, we performed Western blot analysis of representative excitatory postsynaptic markers, including SH3 and multiple ankyrin repeat domains 3 (Shank3), Homer protein homolog 1 (Homer1), and the α-amino-3-hydroxy-5-methyl-4-isoxazolepropionic acid (AMPA) receptor subunit (GluA2). Our results showed a downregulation of these markers in the synaptosomes of both brain regions in *Cyfip2^+/R87C^* mice at PWK 14 and 28 (**Fig 3B**). Interestingly, consistent with the DAVID GO analysis of the DEGs (**Fig 2C**), we observed increased levels of presynaptic markers, such as Synaptotagmin and Synapsin 1, suggesting that synapse remodeling in *Cyfip2^+/R87C^* mice may involve an imbalance between post- and pre-synaptic compartments.

**Fig 3.**
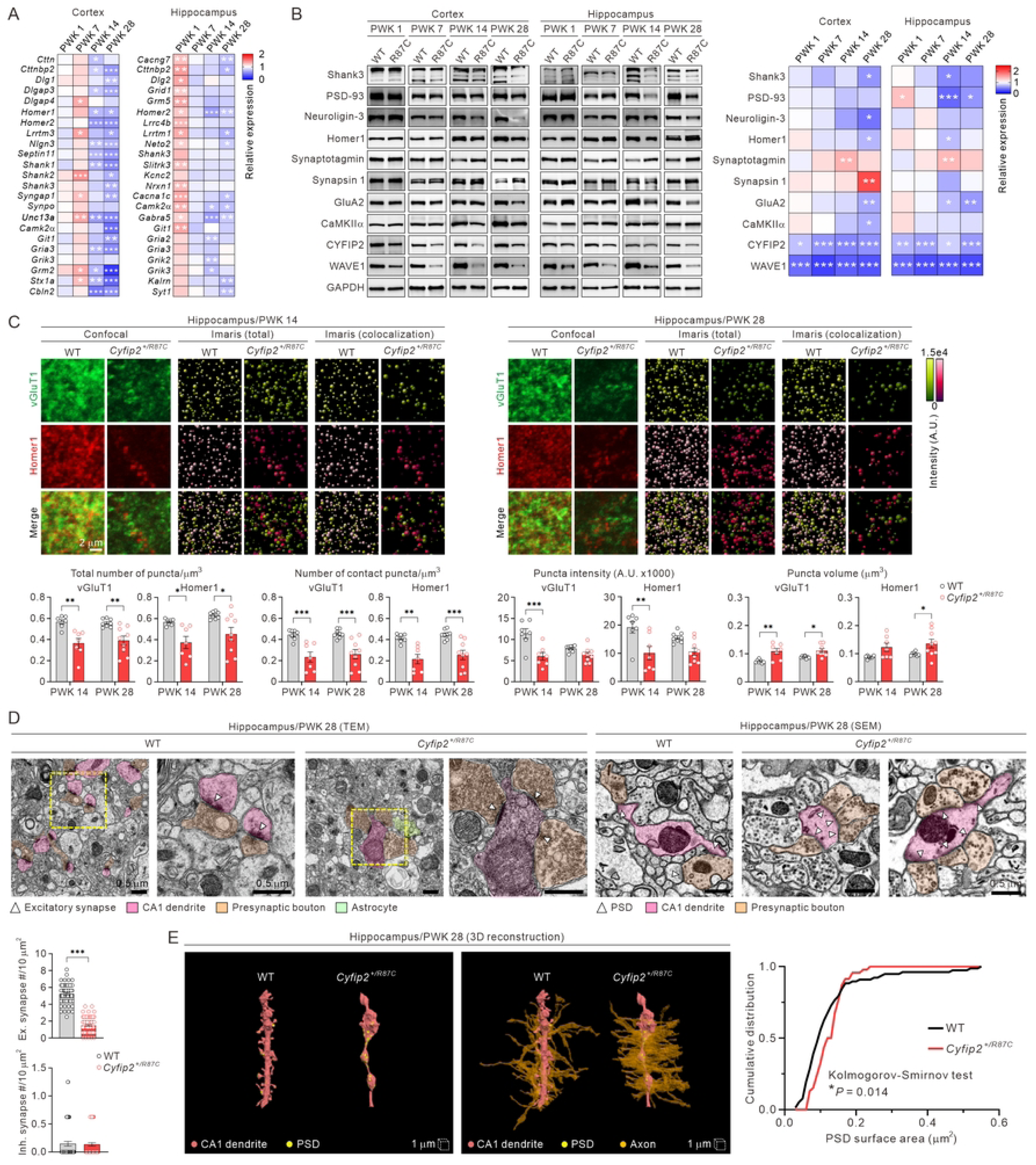
Molecular and structural synaptic remodeling in *Cyfip2^+/R87C^* mice during seizure evolution. (A) Heat maps show qRT-PCR results for DEGs encoding synaptic proteins in the cortex and hippocampus of *Cyfip2^+/R87C^* mice compared to WT mice at various ages (n = 6 mice per genotype, two-way ANOVA with Šídák’s multiple comparisons test). (B) Representative Western blot images and heat maps display the expression levels of representative synaptic marker proteins in the cortical and hippocampal synaptosomes of *Cyfip2^+/R87C^* mice compared to WT mice at different ages (n = 6 to 8 mice per genotype, two-way ANOVA with Šídák’s multiple comparisons test). (C) Representative fluorescence immunohistochemistry images and quantification show changes in excitatory presynaptic (vGluT1) and postsynaptic (Homer1) markers in the hippocampal CA1 region of *Cyfip2^+/R87C^* mice compared to WT mice at PWK 14 and 28 (n = 7 to 9 mice per genotype, two-way ANOVA with Šídák’s multiple comparisons test). Images obtained using Imaris software for automated foci counting are also included. Scale bar, 2 μm. A.U. = arbitrary units. (D) Representative transmission electron microscopy (TEM) and scanning electron microscopy (SEM) images and quantification of excitatory (Ex.) and inhibitory (Inh.) synapse numbers in the hippocampal CA1 region of WT and *Cyfip2^+/R87C^* mice at PWK 28 (n = 59 to 73 images from 3 mice per genotype, unpaired two-tailed Student’s t-test). Presynaptic boutons form synaptic contacts on dendritic spines in WT mice, whereas in *Cyfip2^+/R87C^* mice, they contact dendritic shafts and cluster nearby. PSD = postsynaptic density. Scale bar, 0.5 μm. (E) 3D reconstruction of dendritic segments from CA1 pyramidal neurons in WT and *Cyfip2^+/R87C^* mice (left, without axons; right, with axons). The cumulative graph displays the distributions of PSD surface area in WT and *Cyfip2^+/R87C^* neurons (n = 46 to 77 PSDs per genotype, Kolmogorov-Smirnov test). \**P* < 0.05; \*\**P* < 0.01; \*\*\**P* < 0.001. Data are represented as mean ± standard error of the mean.

Fluorescence immunohistochemistry using representative excitatory presynaptic (vesicular glutamate transporter 1, vGluT1) and postsynaptic (Homer1) markers revealed a decrease in the number of total and contact puncta, as well as a reduction in average puncta intensity in the hippocampal CA1 region of *Cyfip2^+/R87C^* mice compared to WT mice at PWK 14 and 28, indicating a reduced number of excitatory synapses (**Fig 3C and S2 Fig**). Interestingly, the average puncta volume for both vGluT1 and Homer1 increased in *Cyfip2^+/R87C^* mice, suggesting that the remaining synapses may have altered structures. In contrast, vGluT1 and Homer1 parameters in the prelimbic medial prefrontal cortex (mPFC) of *Cyfip2^+/R87C^* mice were largely normal, except for a decrease in Homer1 puncta number and average intensity at PWK 14 (**S3 Fig**).

To further investigate synaptic changes in the hippocampal CA1 region of *Cyfip2^+/R87C^* mice, we conducted ultrastructural analysis using electron microscopy. Consistent with the immunohistochemistry findings, the number of excitatory (asymmetric) synapses decreased, while the number of inhibitory (symmetric) synapses remained normal in *Cyfip2^+/R87C^* mice at PWK 28 (**Fig 3D**). Notably, we observed a significant reduction in dendritic spines, small protrusions on dendrites where excitatory synapses are typically formed, in *Cyfip2^+/R87C^* neurons. Instead, excitatory synapses were predominantly located on dendritic shafts, with multiple presynaptic terminals clustering nearby (**Fig 3D**). This altered synaptic configuration was further highlighted by three-dimensional (3D) reconstruction of dendritic segments from CA1 pyramidal neurons in WT and *Cyfip2^+/R87C^* mice (**Fig 3E**). In *Cyfip2^+/R87C^* neurons, lacking dendritic spines, postsynaptic densities (PSDs) were shown on irregularly shaped dendritic shafts where axons made denser contacts compared to WT neurons. The distribution of PSD surface area differed significantly between WT and *Cyfip2^+/R87C^* neurons, in line with immunohistochemical results showing an increase in synaptic puncta volume. Additionally, *Cyfip2^+/R87C^* axons exhibited excessive myelination, consistent with increased myelin basic protein (MBP) signals in the hippocampus compared to WT mice (**S4 Fig**).

### Gliosis in *Cyfip2^+/R87C^* mice during seizure evolution

Beyond synapse remodeling, transcriptomic analysis of *Cyfip2^+/R87C^* mice revealed a shift towards negative enrichment for neuron-specific gene sets and positive enrichment for glia-specific gene sets as they aged, particularly in the hippocampus (**Fig 2**). To further investigate this at the cellular level, we performed fluorescence immunohistochemistry using antibodies against markers for neurons (neuronal nuclei, NeuN) and various glial cell types (oligodendrocyte transcription factor 2 (Olig2) for oligodendrocytes, ionized calcium-binding adapter molecule 1 (IBA1) for microglia, and glial fibrillary acidic protein (GFAP) for astrocytes).

In the prelimbic mPFC, NeuN mean intensity in *Cyfip2^+/R87C^* mice was elevated compared to WT mice at PWK 1 but became comparable at later stages (**S5A Fig**). Interestingly, glial markers in *Cyfip2^+/R87C^* mice showed a sequential increase with age. Specifically, Olig2 mean intensity increased at PWK 7, followed by IBA1 at PWK 14, and GFAP at both PWK 14 and 28, compared to WT mice. In the mPFC of *Cyfip2^+/-^* mice at PWK 28, Olig2 mean intensity increased on the C57BL/6N background, and GFAP mean intensity increased on the C57BL/6J background, relative to their respective WT mice (**S5A Fig**).

In the hippocampal CA1 region of *Cyfip2^+/R87C^* mice, NeuN mean intensity decreased by 50% compared to WT mice at PWK 14 and 28, coinciding with the onset of seizures and lethality in these mice (**Fig 4**). A closer examination of the region revealed dispersion of CA1 pyramidal neurons in *Cyfip2^+/R87C^* mice. Similar to the mPFC, glial markers in the hippocampal CA1 region of *Cyfip2^+/R87C^* mice exhibited an age-related increase compared to WT mice, with oligodendrocyte, microglia, and astrocytes in that order (**Fig 4**), consistent with the results of GSEA of the transcriptome for cell-type-specific gene sets (**Fig 2D**). However, unlike the mPFC, Olig2 mean intensity in the hippocampus increased at PWK 7 and persisted through PWK 14 and 28. In the hippocampus of *Cyfip2^+/-^* mice at PWK 28, GFAP mean intensity, but not other markers, was mildly increased compared to WT mice on both the C57BL/6N and C57BL/6J backgrounds, although the increase was much less pronounced than in *Cyfip2^+/R87C^* mice (**S5B Fig**).

**Fig 4.**
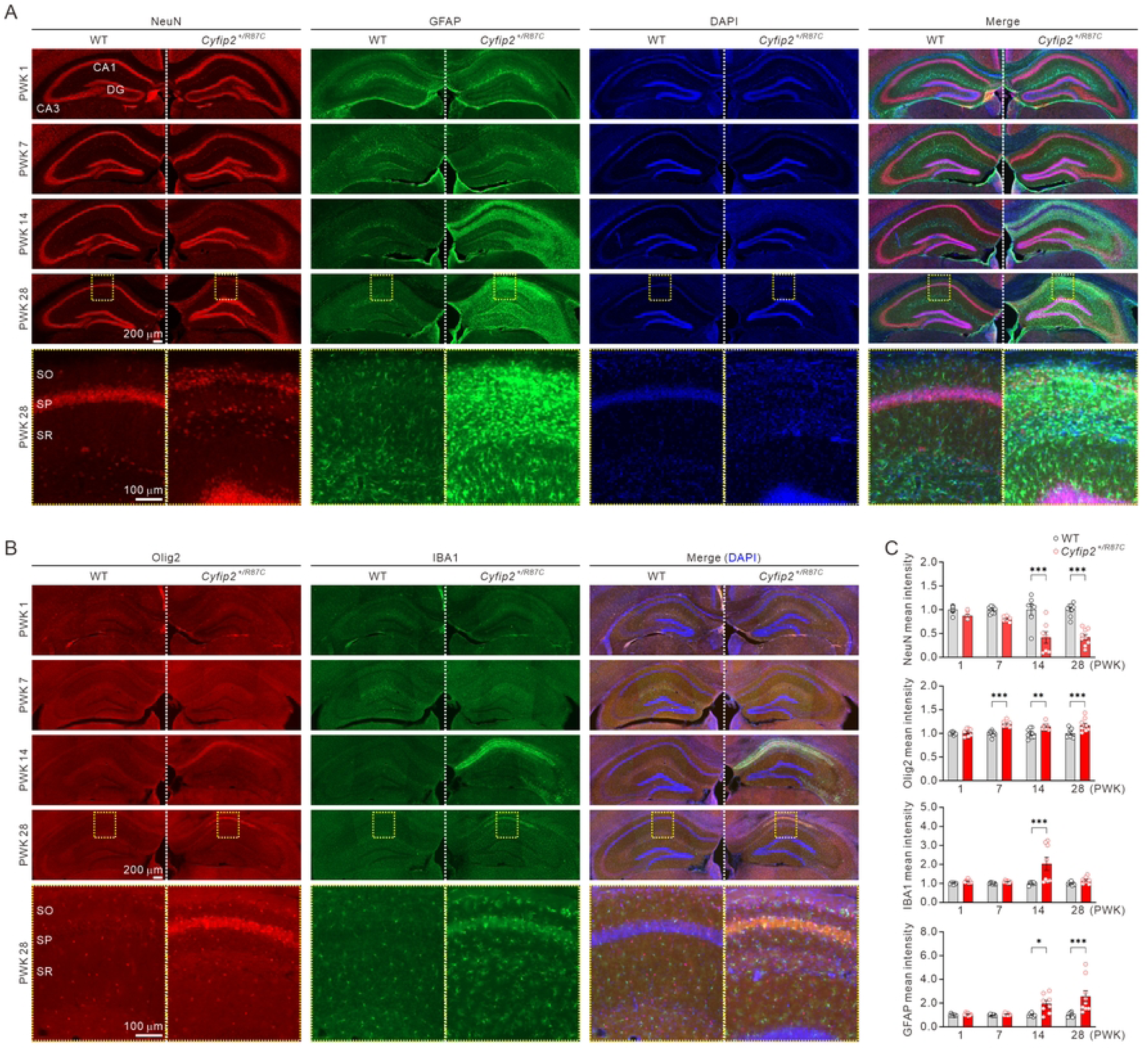
Neuronal and glial changes in the hippocampus of *Cyfip2^+/R87C^* mice during seizure evolution. (A) Representative fluorescence immunohistochemistry images illustrate age-dependent changes in the mean intensities of neuronal (NeuN) and astrocytic (GFAP) markers in the hippocampus of *Cyfip2^+/R87C^* mice compared to WT mice. DAPI was used for counterstaining nuclei. High-magnification images of the CA1 region at PWK 28 are shown. SO = stratum oriens, SP = stratum pyramidale, SR = stratum radiatum. (B) Representative fluorescence immunohistochemistry images show age-dependent changes in the mean intensities of oligodendrocytic (Olig2) and microglial (IBA1) markers in the hippocampus of *Cyfip2^+/R87C^* mice compared to WT mice. (C) Bar graphs display relative changes in mean intensities of neuronal and glial markers in the hippocampal CA1 region of *Cyfip2^+/R87C^* mice compared to age-matched WT mice (n = 7 to 9 mice per genotype, two-way ANOVA with Šídák’s multiple comparisons test). \**P* < 0.05; \*\**P* < 0.01; \*\*\**P* < 0.001. Data are represented as mean ± standard error of the mean.

These results suggest that neurons and multiple glial cell types undergo dynamic changes in *Cyfip2^+/R87C^* mice during seizure evolution. Notably, the sequential nature of these phenomena indicates that changes in neurons and various glial cells may influence each other. Thus, in the next phase of our study, we focused on microglia and astrocytes to investigate these specific changes in greater detail.

### Changes in microglial morphology and activity in *Cyfip2^+/R87C^* mice

We analyzed the 3D morphology of microglia in the prelimbic mPFC and CA1 hippocampus of WT and *Cyfip2^+/R87C^* mice at PWK 14 and 28. These stages were selected as they correspond to the onset of lethality and seizures in *Cyfip2^+/R87C^* mice and because IBA1 intensity increased at PWK 14 and normalized by PWK 28 in *Cyfip2^+/R87C^* brains (**Fig 4**). Understanding these changes may provide insights into the roles of microglia and their interactions with other cell types during seizure evolution.

In the mPFC, Sholl analysis showed decreased intersections at certain distances for microglia in *Cyfip2^+/R87C^* mice compared to WT mice at PWK 14, and slightly increased intersections at some distances at PWK 28 (**S6 Fig**). However, average values for total branch length, area, branch points, and maximal diameter of microglia were comparable between genotypes at both ages. Morphological changes in microglia were more pronounced in the hippocampus of *Cyfip2^+/R87C^* mice (**Fig 5A**). Sholl analysis revealed significantly fewer intersections at most distances in *Cyfip2^+/R87C^* hippocampus compared to WT at both PWK 14 and 28. Additionally, total branch length and area were reduced in *Cyfip2^+/^*^R87C^ microglia at both ages. At PWK 28, but not PWK 14, the number of total branch points decreased, while the maximal branch diameter increased in *Cyfip2^+/R87C^* microglia. These morphological properties suggest the presence of “ameboid microglia” in the hippocampus of *Cyfip2^+/R87C^* mice, which are highly active (i.e., phagocytic) and often associated with pathological brain conditions [26].

**Fig 5.**
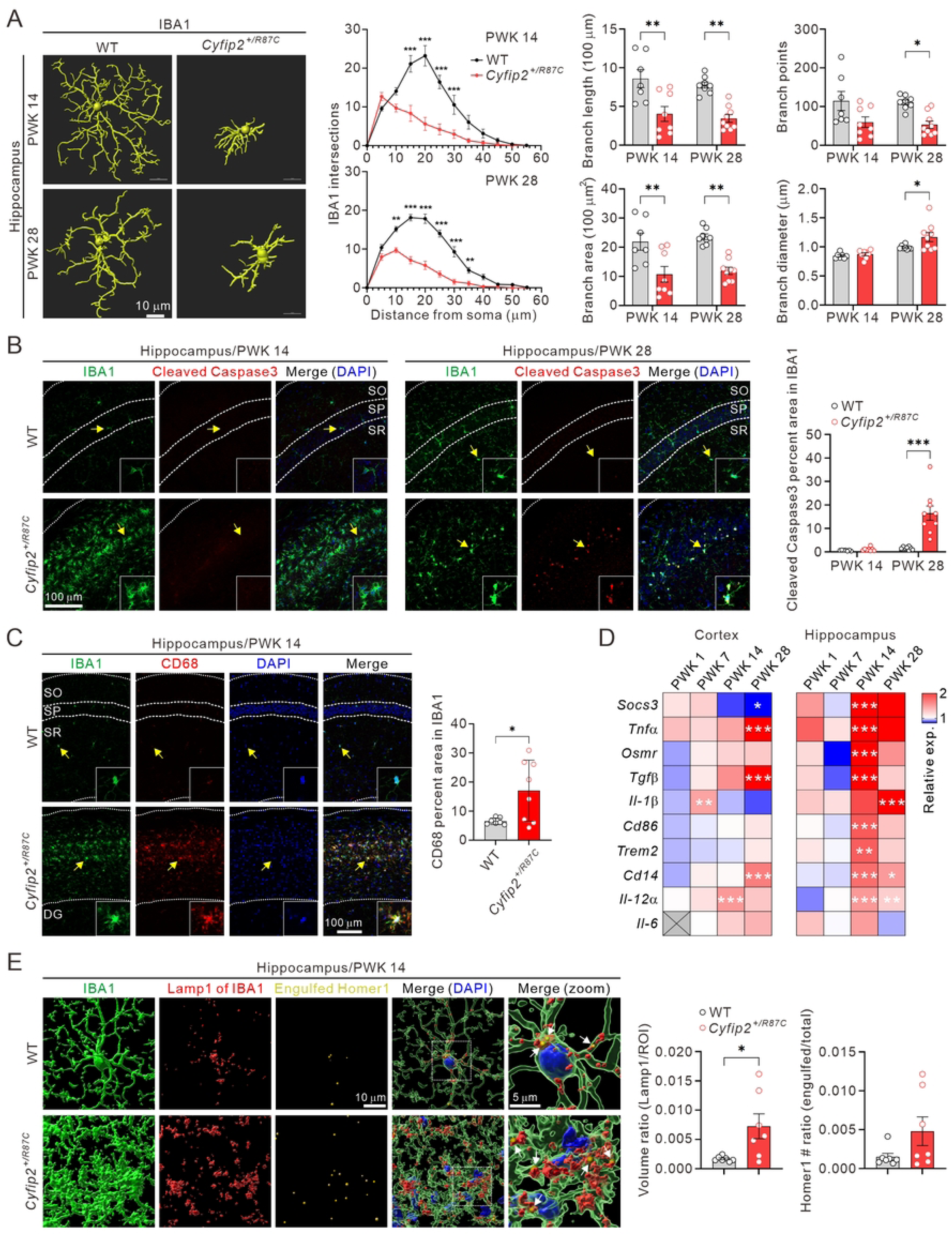
Morphological and functional changes of microglia in *Cyfip2^+/R87C^* mice. (A) Representative 3D images of microglia in the hippocampal CA1 region of WT and *Cyfip2^+/R87C^* mice at PWK 14 and 28, processed using Imaris software. Graphs show quantifications from Sholl analysis as well as measurements of branch length, area, points, and diameter (n = 7 to 9 mice per genotype, two-way ANOVA with Šídák’s multiple comparisons test). (B) Representative fluorescence immunohistochemistry images and quantification show increased cleaved Caspase3 signals in microglia of the *Cyfip2^+/R87C^* hippocampus compared to the WT hippocampus at PWK 28, but not at PWK 14 (n = 7 to 9 mice per genotype, two-way ANOVA with Šídák’s multiple comparisons test). Arrows indicate microglia with enlarged images shown in the insets. SO = stratum oriens, SP = stratum pyramidale, SR = stratum radiatum. (C) Representative fluorescence immunohistochemistry images and quantification show increased CD68 signals in microglia of the *Cyfip2^+/R87C^* hippocampus compared to the WT hippocampus at PWK 14 (n = 7 to 8 mice per genotype, unpaired two-tailed Student’s t-test). (D) Heat maps display qRT-PCR results for genes related to microglial activation in the cortex and hippocampus of *Cyfip2^+/R87C^* mice compared to WT mice at different ages (n = 6 mice per genotype, two-way ANOVA with Šídák’s multiple comparisons test). Note that *Il-6* was not detected in the cortex at PWK 1. (E) Representative 3D images illustrate microglia (IBA1), lysosome (Lamp1) within the microglia, and excitatory synaptic protein (Homer1) within microglial lysosome (i.e., engulfed Homer1, indicated by white arrows in merged images) in the hippocampal CA1 region of WT and *Cyfip2^+/R87C^* mice at PWK 14, analyzed using Imaris software. Graphs show quantifications of microglial lysosome and engulfed Homer1 in WT and *Cyfip2^+/R87C^* mice (n = 7 mice per genotype, unpaired two-tailed Student’s t-test). \**P* < 0.05; \*\**P* < 0.01; \*\*\**P* < 0.001. Data are represented as mean ± standard error of the mean.

To explore whether the normalization of IBA1 intensity at PWK 28 was related to microglial death, we investigated apoptotic markers. Cleaved Caspase3 signals in microglia were significantly higher in the *Cyfip2^+/R87C^* hippocampus at PWK 28, but not PWK 14, compared to WT, suggesting increased apoptotic cell death at this stage (**Fig 5B**). Consistent with the observed morphological changes, CD68 signals, a marker for activated microglia, were significantly higher in the hippocampus of *Cyfip2^+/R87C^* mice compared to WT mice (**Fig 5C**). Additionally, expression levels of multiple genes associated with microglial activation, such as inflammatory cytokines, significantly increased in *Cyfip2^+/R87C^* mice, particularly in the hippocampus at PWK 14 (**Fig 5D**).

Lastly, we examined whether activated microglia contributed to the reduction of excitatory synapses in the hippocampal CA1 region of *Cyfip2^+/R87C^* mice at PWK 14 through synapse engulfment [27]. We observed an increase in the total volume of lysosomes (labeled by Lamp1) within microglia in the *Cyfip2^+/R87C^* hippocampus compared to WT (**Fig 5E**). Although the amount of Homer1 within microglial lysosomes, representing engulfed excitatory synapses, showed a trend towards increase, it did not reach statistical significance. This may be due to other mechanisms contributing to synapse reduction or because microglial phagocytosis of synapses occurred earlier than PWK 14, by which time the number of excitatory synapses had already significantly decreased (**Fig 3**).

### Lipid droplets and crystals in astrocytes of *Cyfip2^+/R87C^* mice

We next investigated astrocytes, marked by GFAP, which showed increased intensity in the mPFC and hippocampus of *Cyfip2^+/R87C^* mice at both PWK 14 and 28 (**Fig 4**). In the prelimbic mPFC, astrocytes from *Cyfip2^+/R87C^* mice exhibited increased intersections in Sholl analysis, although no significant differences in other branch parameters were observed compared to WT mice (**S7 Fig**). In the hippocampus, however, Sholl analysis revealed decreased intersections at most distance points for *Cyfip2^+/R87C^* astrocytes relative to WT astrocytes at both PWK 14 and 28 (**Fig 6A**). Additionally, average branch length, area, and points were reduced, while branch diameter increased in *Cyfip2^+/R87C^* astrocytes at both ages.

**Fig 6.**
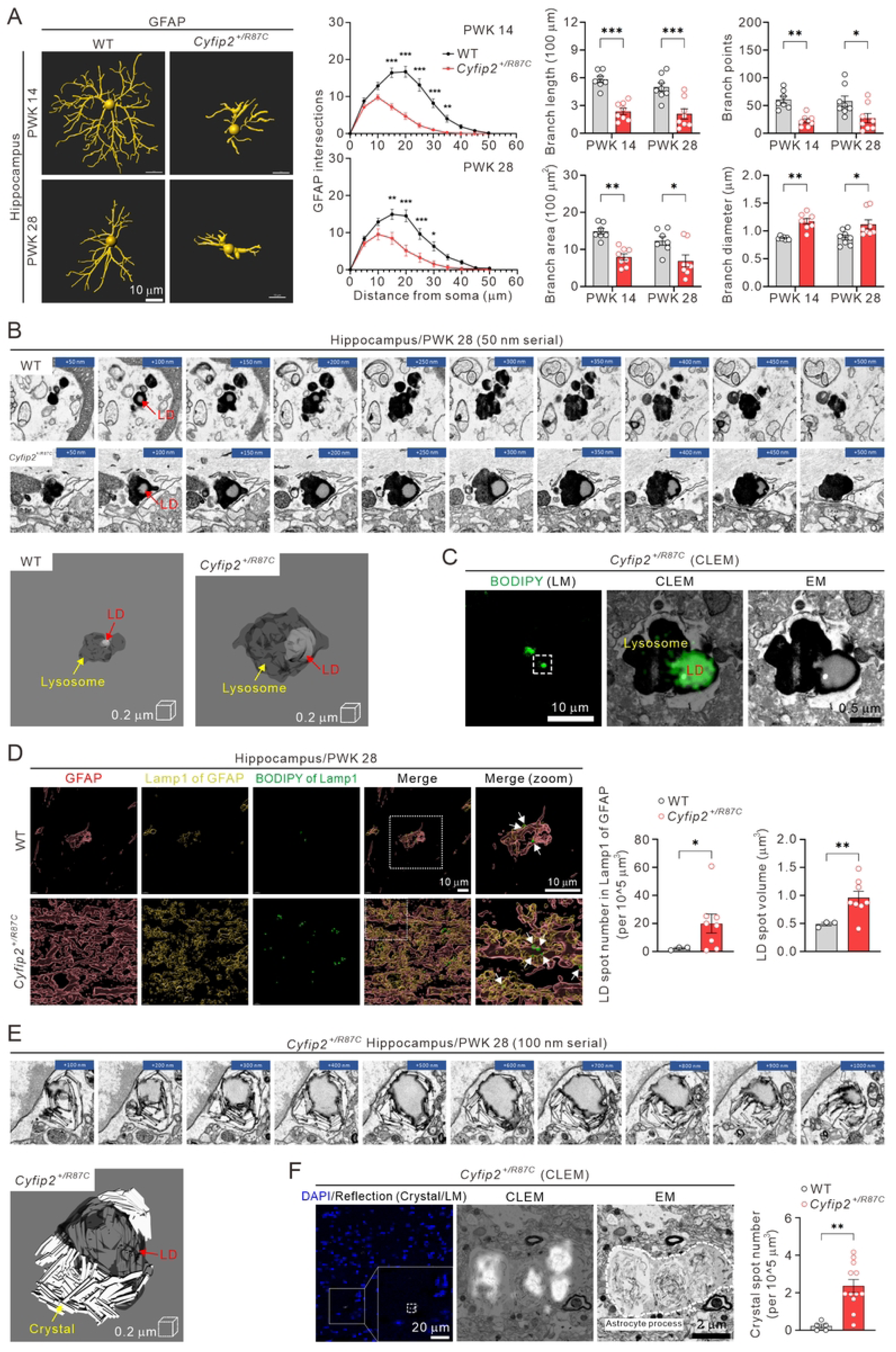
Changes in morphology and lipid droplets of astrocytes in *Cyfip2^+/R87C^* mice. (A) Representative 3D images of astrocytes in the hippocampal CA1 region of WT and *Cyfip2^+/R87C^* mice at PWK 14 and 28, processed using Imaris software. Graphs show quantifications from Sholl analysis as well as measurements of branch length, area, points, and diameter (n = 7 to 8 mice per genotype, two-way ANOVA with Šídák’s multiple comparisons test). (B) Serial electron microscopic images of lysosomes containing lipid droplets (LDs) in the hippocampal astrocytes of WT and *Cyfip2^+/R87C^* mice at PWK 28. 3D-rendered images of these lysosomes are also shown. (C) Correlative light and electron microscopy (CLEM) image of the hippocampal astrocyte in *Cyfip2^+/R87C^* mouse at PWK 28 shows BODIPY (light microscopy, LM), a marker for neutral lipids, within electron-dense lysosome observed by electron microscopy (EM). (D) Representative 3D images illustrate astrocytes (GFAP), lysosome (Lamp1) within the astrocytes, and LDs (BODIPY) within astrocytic lysosome (indicated by white arrows in merged images) in the hippocampus of WT and *Cyfip2^+/R87C^* mice at PWK 28, analyzed using Imaris software. Graphs show quantifications of the number and volume of LDs within astrocytic lysosomes in WT and *Cyfip2^+/R87C^* mice (n = 3 to 8 mice per genotype, unpaired two-tailed Student’s t-test). (E) Serial electron microscopic images of a lipid droplet with crystals in the hippocampal astrocyte of *Cyfip2^+/R87C^* mouse at PWK 28. A 3D-rendered image of this lipid droplet is also presented. (F) CLEM image of the hippocampal astrocyte in *Cyfip2^+/R87C^* mouse at PWK 28 shows reflection signals from LM that match the crystals observed by EM. The graph displays the number of crystal spots, as measured by reflection signals from LM, in the hippocampus of WT and *Cyfip2^+/R87C^* mice at PWK 28 (n = 5 to 12 mice per genotype, unpaired two-tailed Student’s t-test). \**P* < 0.05; \*\**P* < 0.01; \*\*\**P* < 0.001. Data are represented as mean ± standard error of the mean.

Given the persistent increase in GFAP intensity and morphological changes in *Cyfip2^+/R87C^* mice, coupled with recurrent seizures, we hypothesized that additional features in *Cyfip2^+/R87C^* astrocytes might be associated with these seizures. Using electron microscopy, we discovered unexpected alterations in the lysosomes of *Cyfip2^+/R87C^* astrocytes in the hippocampus at PWK 28 (**Fig 6B**). Specifically, lysosomes in *Cyfip2^+/R87C^* astrocytes contained larger lipid droplets (LDs) compared to those in WT astrocytes. Correlative light and electron microscopy (CLEM) with BODIPY, a marker for neutral lipids, confirmed the presence of LDs in the lysosomes (**Fig 6C**). Quantitative analysis through fluorescence immunohistochemistry revealed that both the number and volume of LDs within astrocytic lysosomes were significantly increased in the hippocampus of *Cyfip2^+/R87C^* mice compared to WT mice at PWK 28, but not at PWK 14 (**Fig 6D and S8 Fig**).

While examining electron microscopy images, we frequently observed crystals on or within the LDs of astrocytes in the hippocampus of *Cyfip2^+/R87C^* mice, but never in WT mice (**Fig 6E**). Although the exact identity of these crystals remains unknown, cholesterol crystallization is well-documented in hepatocyte LDs in nonalcoholic steatohepatitis (NASH) and on LD surfaces in atherosclerotic lesions [28, 29]. Using CLEM, we confirmed that these crystals could be visualized through reflection imaging [30], and quantification showed a higher number of crystal spots in the hippocampus of *Cyfip2^+/R87C^* mice compared to WT mice (**Fig 6F**). Notably, we identified various types of LDs, ranging from those without crystallization to those with moderate and extensive crystallization, which may represent different stages of lipid accumulation or processing (**S9 Fig**).

### Single-nucleus transcriptomic analysis in the *Cyfip2^+/R87C^* hippocampus

Although we observed changes in LDs mainly in astrocytes, not in neurons or other glial cell types of *Cyfip2^+/R87C^* mice, we hypothesized that lipid-related pathways might also be affected in other cell types, as lipids are transported in various forms among neurons and glial cells to maintain proper lipid metabolism and storage in the brain [31]. Furthermore, we aimed to gain a deeper understanding of the molecular characteristics of astrocytes in *Cyfip2^+/R87C^* mice, particularly focusing on whether and how they influence neuronal function and contribute to seizures. To test this hypothesis in a cell-type-specific manner, we conducted single-nucleus transcriptomic analysis on the hippocampus of WT and *Cyfip2^+/R87C^* mice at P28 WK.

After applying stringent quality controls, including doublet removal, we retained 54,998 high-quality nuclei for downstream analyses (**S10A Fig**). Uniform Manifold Approximation and Projection (UMAP) visualization identified twelve distinct cell populations (**Fig 7A**), including two major neuronal groups (excitatory and inhibitory), five glial lineages (astrocytes, committed oligodendrocyte precursor cells (OPCs), OPCs, oligodendrocytes, and microglia), a rare neuronal subtype (Cajal-Retzius), a fibroblast population, endothelial and ependymal cells, and a “mixed” cluster co-expressing markers characteristic of both neurons and oligodendrocytes (**Fig 7B and S4 Table**). While some subsets of neurons and oligodendrocytes exhibited partial segregation by genotype, the substantial intermixing of replicates within each genotype indicated minimal batch effects, making batch correction unnecessary (**S10B Fig**).

**Fig 7.**
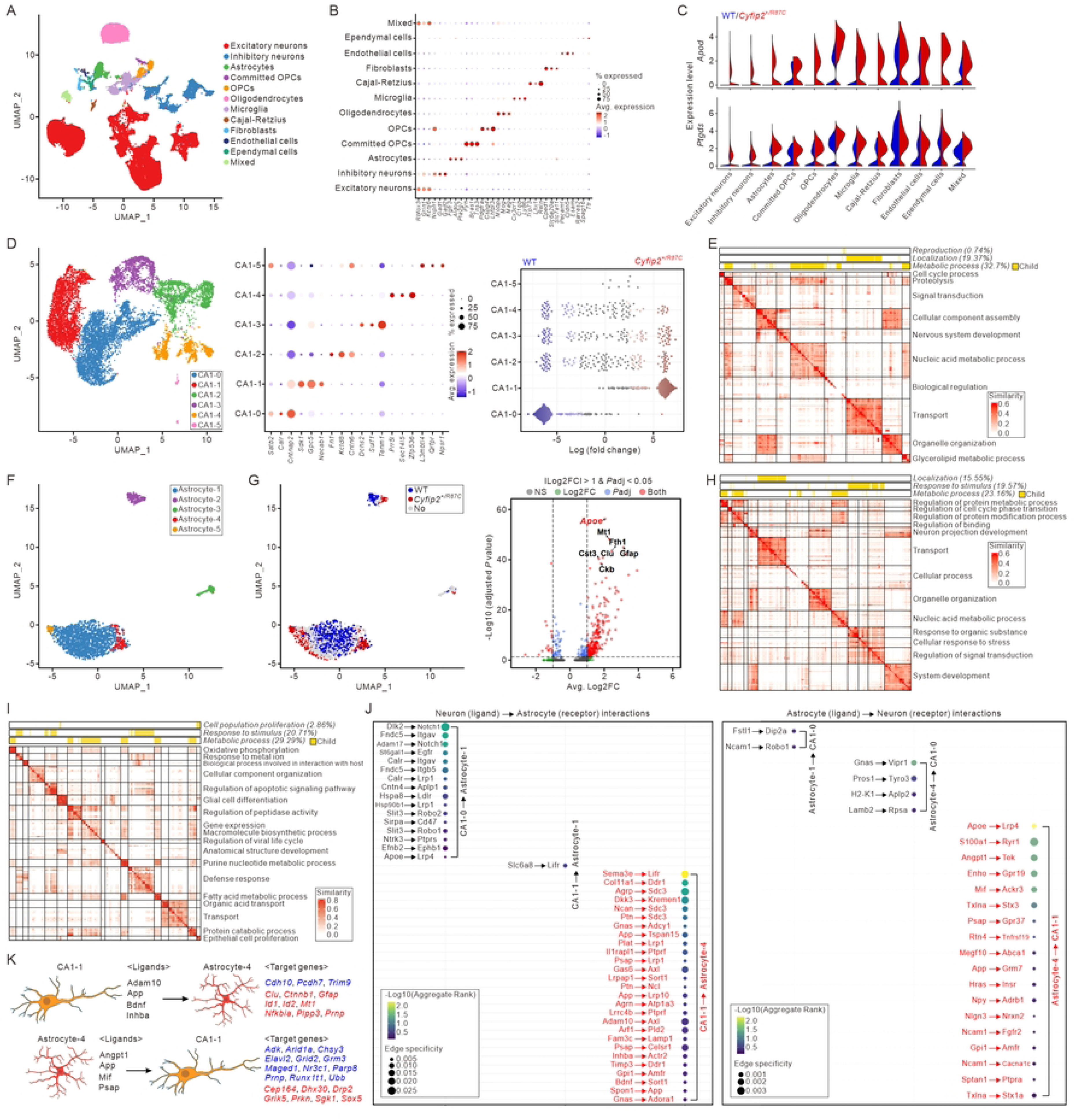
Single-nucleus transcriptomic analysis on the hippocampus of *Cyfip2^+/R87C^* mice. (A) UMAP visualization of twelve hippocampal cell types from WT and *Cyfip2^+/R87C^* mice at PWK 28. (B) A dot plot illustrating the expression of cell-specific genes used to identify the twelve cell types, with dot size representing the percentage of cells expressing the gene and color indicating the average (Avg.) expression level. (C) Violin plot showing the expression levels of *Apod* and *Ptgds* in different cell types, grouped by genotype. (D) Subclustering results of CA1 neurons. The left panel presents a UMAP visualization of CA1 neuronal subtypes, while the middle panel features a dot plot highlighting the top DEGs for each cluster. The right panel shows the results of MiloR analysis, illustrating CA1 abundance differences, with genotypes color-coded as WT (blue) and *Cyfip2^+/R87C^* (red). (E) Heatmap displaying Over-Representation Analysis (ORA) results for regulons regulated by transcription factors (TFs) with higher activity in CA1-1 compared to CA1-0. Color intensity represents similarity, and each label corresponds to the ancestor term of the respective cluster. (F) Subclustering results of astrocytes. (G) The left panel shows a UMAP visualization of abundance analysis results, illustrating differences in astrocyte abundance between WT (blue) and *Cyfip2^+/R87C^* (red) mice. The right panel displays a volcano plot illustrating the results of the differential expression (DE) test between Astrocyte-4 and other subtypes. (H) Heatmap illustrating the ORA results for regulons governed by TFs with higher activity in Astrocyte-4 compared to Astrocyte-1. (I) Heatmap showing GSEA results for Astrocyte-4 compared to Astrocyte-1. (J) Ligand-receptor (LR) interactions between subtypes of neurons and astrocytes. Interactions were inferred from the expression profiles of ligands and receptors across different cell populations. LR interactions specific to *Cyfip2^+/R87C^* neuron-astrocyte pairs (CA1-1 to Astrocyte-4 and Astrocyte-4 to CA1-1) are highlighted in red. (K) Diagrams depicting bidirectional interactions between CA1-1 and Astrocyte-4, illustrating the potential influence of ligands on target cell gene expression (blue, downregulated; red, upregulated).

Notably, a global differential expression (DE) analysis revealed significant upregulation of genes involved in lipid metabolism and transport in the *Cyfip2^+/R87C^* condition, with apolipoprotein D (*Apod*) and prostaglandin D2 synthase (*Ptgds*) being particularly prominent (**Fig 7C and S10C Fig and S5 Table**). Focusing on the neuronal compartment, initial sub-clustering separated excitatory neurons from inhibitory neurons (**S10D Fig**). The excitatory neurons were further divided into subgroups, including pyramidal, granule, and mossy cells (**S10E Fig**). Among the excitatory neurons, pyramidal neurons were classified into five distinct subtypes, such as canonical CA1 and CA2 populations, defined by specific marker genes and functional annotations (**S10F Fig**). Further sub-clustering of CA1 neurons identified six distinct subsets, with notable differences in abundance between the CA1-0 (WT) and CA1-1 (*Cyfip2^+/R87C^*) subsets (**Fig 7D**). Regulatory network analysis using Single-Cell rEgulatory Network Inference and Clustering (SCENIC) [32] revealed transcription factors with significantly increased regulon activity in CA1-1 compared to CA1-0. These transcription factors are associated with processes such as cellular metabolism and transport (**Fig 7E**). Additionally, GSEA of DEGs in CA1-1 neurons demonstrated enrichment for catabolic and metabolic processes, including phosphatidylinositol dephosphorylation (**S10G Fig**).

Among the glial populations, oligodendrocytes were divided into two distinct subtypes, one of which was notably characterized by elevated expression of lipid-associated genes, including *Apod* and *Ptgds* (**S11A Fig**). Microglia were categorized into six condition-dependent subtypes (**S11B Fig**); however, neither oligodendrocytes nor microglia showed significant genotype-dependent differences in their relative abundance. In contrast, astrocytes, which were divided into five subtypes (**Fig 7F**), exhibited more pronounced genotype-specific changes. Notably, Astrocyte-4, which was more abundant in *Cyfip2^+/R87C^* samples, showed a significant increase in the expression of apolipoprotein E (*Apoe*), a lipid-associated gene, and *Gfap* compared to the other astrocyte subtypes (**Fig 7G**). SCENIC analysis identified transcription factors with significantly elevated regulon activity in Astrocyte-4 compared to Astrocyte-1, regulating gene networks linked to cellular metabolism, stimulus response, and transport processes (**Fig 7H**). Complementing these findings, GSEA of Astrocyte-4 highlighted metabolic pathways, including oxidative phosphorylation, fatty acid metabolism, and related processes (**Fig 7I**).

Using gene expression profiles, we analyzed ligand-receptor (LR) interactions between neurons and astrocytes to investigate their potential influence on each other in *Cyfip2^+/R87C^* mice. Notably, neuron-astrocyte pairs specific to *Cyfip2^+/R87C^* mice (i.e., CA1-1 to Astrocyte-4 and Astrocyte-4 to CA1-1) exhibited 27 and 18 unique LR interactions, respectively, compared to control neuron-astrocyte pairs (e.g., CA1-0 to Astrocyte-1 and Astrocyte-1 to CA1-0 in WT mice) (**Fig 7J**). Among the 27 CA1-1 to Astrocyte-4 LR interactions, a disintegrin and metalloproteinase domain 10 (Adam10), amyloid-beta precursor protein (App), brain derived neurotrophic factor (Bdnf), and inhibin subunit beta A (Inhba) acting as ligands on CA1-1, were predicted to have significant regulatory potential, influencing twelve target genes in Astrocyte-4, including *Gfap*, prion protein (*Prnp*), phospholipid phosphatase 3 (*Plpp3*), and protocadherin 7 (*Pcdh7*) (**S12A Fig**) [33]. Notably, *Gfap*, *Prnp*, and *Plpp3* were significantly upregulated, whereas *Pcdh7* was downregulated, in Astrocyte-4 compared to Astrocyte-1 (**Fig 7K and S12B Fig**). In the opposite LR direction, among the 18 Astrocyte-4 to CA1-1 LR interactions, four ligands (angiopoietin 1 (Angpt1), App, macrophage migration inhibitory factor (Mif), and prosaposin (Psap)) on Astrocyte-4 were predicted to have significant regulatory potential, influencing a total of 28 target genes in CA1-1 (**S12C Fig**). These 28 genes were most significantly associated with glutamatergic synaptic transmission (**S12D Fig**). Of these, 19 target genes, including adenosine kinase (*Adk*, downregulated) and glutamate ionotropic receptor kainate type subunit 5 (*Grik5*, upregulated), were significantly altered in CA1-1 compared to CA1-0 (**Fig 7K and S12E Fig**).

### Brain proteomic and lipidomic changes in *Cyfip2^+/R87C^* mice

The single-nucleus transcriptomic analysis suggested alterations in lipid and metabolic homeostasis, as well as changes in intercellular crosstalk in the *Cyfip2^+/R87C^* brain, prompting further investigation into the proteomic and lipidomic profiles of these mice. We performed proteomic analysis on the hippocampus of WT and *Cyfip2^+/R87C^* mice at both PWK 14 and 28. Principal component analysis (PCA) revealed a distinct separation in proteomic profiles between WT and *Cyfip2^+/R87C^* mice at both ages (**Fig 8A**). A total of 3,102 and 3,748 differentially expressed proteins (DEPs) were identified in the hippocampus of *Cyfip2^+/R87C^* mice compared to WT mice at PWK 14 and 28, respectively (**Fig 8B and S6 Table**). Notably, an increased abundance of Apod and Ptgds was confirmed in *Cyfip2^+/R87C^* mice relative to WT mice, particularly at PWK 28 (**Fig 8C**). Clustering analysis of the entire proteome separated them into four (A to D) groups based on age-dependent relative expression levels in *Cyfip2^+/R87C^* mice compared to WT mice (**Fig 8D and S7 Table**). We focused on groups B and D, which represent proteins that exhibited further decreases and increases, respectively, in *Cyfip2^+/R87C^* mice compared to WT mice as they aged. We hypothesized that these proteins might be associated with the age-dependent seizure severity observed in *Cyfip2^+/R87C^* mice.

**Fig 8.**
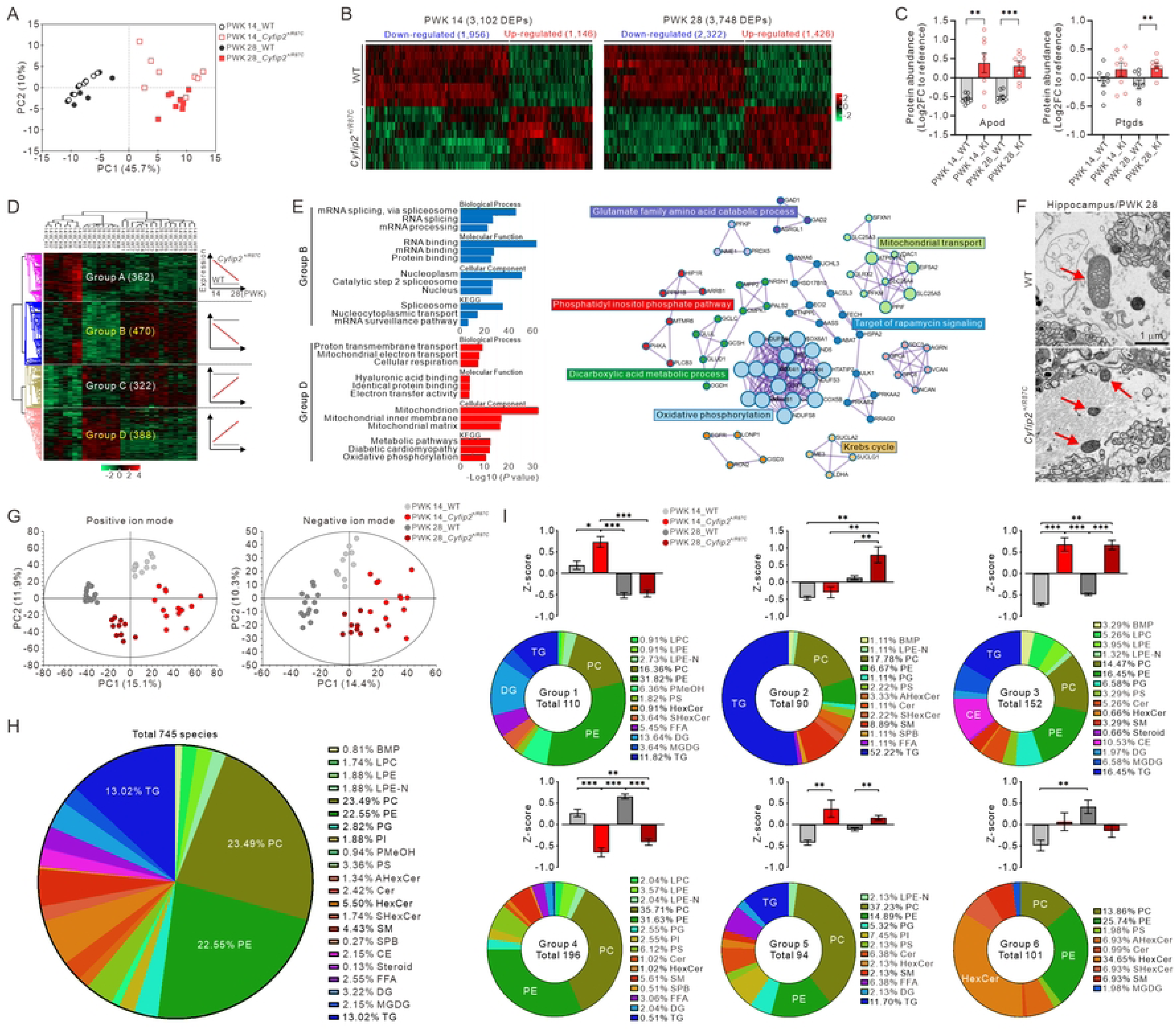
Proteomic and lipidomic changes in the hippocampus of *Cyfip2^+/R87C^* mice. (A) Principal component analysis (PCA) score plots show distinct separations in the proteome between WT and *Cyfip2^+/R87C^* mice at PWK 14 and 28. (B) Heatmaps display 3,102 and 3,748 differentially expressed proteins (DEPs) in the hippocampus of *Cyfip2^+/R87C^* mice at PWK 14 and 28, respectively. (C) Graphs depict the abundance of Apod and Ptgds proteins in WT and *Cyfip2^+/R87C^* mice (n = 8 mice per genotype, Welch’s t-test). KI, knock-in. (D) Heatmap illustrates the separation of four distinct protein groups in the proteome of WT and *Cyfip2^+/R87C^* mice, categorized based on age-dependent relative abundance differences between the genotypes. (E) GO analysis of proteins in groups B and D (left panel). Protein-protein interaction (PPI) networks for group D proteins, highlighting their associated metabolic processes (right panel). (F) Representative electron microscopy images show astrocytic mitochondria (indicated by red arrows) in the hippocampus of WT and *Cyfip2^+/R87C^* mice at PWK 28. (G) PCA score plots display clear separations in the lipidome between WT and *Cyfip2^+/R87C^* mice at PWK 14 and 28. (H) A pie chart illustrates the distribution of identified lipid classes in the hippocampus. (I) Pie charts depict the distribution of identified lipid classes across the six groups classified based on the age-dependent relative expression levels in *Cyfip2^+/R87C^* mice compared to WT mice. Bar graphs present Z-scores for the four combinations (two ages and two genotypes) across the six groups (n = 9 to 14 mice per genotype, Mann-Whitney test).

GO analysis of group B revealed significant enrichment in mRNA splicing and processing (**Fig 8E and S13A Fig**), which is consistent with the downregulation of genes involved in these pathways observed in bulk-tissue transcriptomic analyses from the cortex and hippocampus of *Cyfip2^+/R87C^* mice at PWK 28 (**Fig 2**). Additionally, protein-protein interaction (PPI) networks of group B proteins highlighted processes related to mRNA splicing, processing, and transport (**S14 Fig**). To further explore these findings, we examined alternative splicing events (ASEs) in the cortex and hippocampus of *Cyfip2^+/R87C^* mice compared to WT mice using bulk-tissue transcriptome data. In line with the progressive downregulation of genes involved in mRNA splicing and processing in *Cyfip2^+/R87C^* mice, only three ASE changes were observed at PWK 1 (two in the cortex and one in the hippocampus), and none at PWK 7 (**S8 Table**). However, at PWK 14, we identified three significant ASE changes in the cortex and 15 in the hippocampus, predominantly involving exon skipping, in *Cyfip2^+/R87C^* mice compared to WT mice. Notably, altered exon skipping in the Neurexin 2 (*Nrxn2*) gene, which encodes a transmembrane protein regulating excitatory synapse formation [34], was consistently detected in both the cortex and hippocampus of *Cyfip2^+/R87C^* mice. At PWK 28, we observed 26 significant ASE changes in the cortex and 21 in the hippocampus of *Cyfip2^+/R87C^* mice compared to WT mice. Altered exon skipping in *Nrxn3*, another member of *Nrxn* gene family [35], was identified in the cortex but not in the hippocampus. Additionally, changes in U2 small nuclear RNA auxiliary factor 1 (*U2af1*), spermine oxidase (*Smox*), and phosphoseryl-tRNA kinase (*Pstk*) genes were detected in both brain regions of *Cyfip2^+/R87C^* mice.

GO analysis of group D proteins revealed enrichment in mitochondrial functions, including oxidative phosphorylation and electron transport pathways (**Fig 8E and S13B Fig**). Additionally, PPI networks of group D proteins highlighted several mitochondria-associated metabolic processes (**Fig 8E**). To validate these proteomic changes, we revisited the electron microscopic images of astrocytes in *Cyfip2^+/R87C^* mice at PWK 28. Intriguingly, astrocytic mitochondria in *Cyfip2^+/R87C^* mice were smaller and lacked distinct cristae structures compared to those in WT mice (**Fig 8F**), a mitochondrial characteristic often linked to aging and disease states [36].

We conducted lipidomic analysis on the hippocampus of WT and *Cyfip2^+/R87C^* mice at both PWK 14 and 28. PCA revealed distinct separations in lipidomic profiles based on genotype and age (**Fig 8G**). A total of 745 lipid species were identified, with phosphatidylethanolamine (PE) and phosphatidylcholine (PC) being the most abundant lipid classes (**Fig 8H**), consistent with recent reports on the mouse brain lipidome [37]. Similar to the proteomic analysis, clustering analysis of the entire lipidome categorized them into six groups based on age-dependent relative expression levels in *Cyfip2^+/R87C^* mice compared to WT mice (**S15 Fig and S9 Table**). Groups 1 to 3 represented lipid classes upregulated in *Cyfip2^+/R87C^* mice compared to WT mice (**Fig 8I**). Specifically, group 1 lipids were upregulated at PWK 14 in *Cyfip2^+/R87C^* mice compared to WT mice at both PWK 14 and 28, as well as compared to *Cyfip2^+/R87C^* mice at PWK 28. Group 2 lipids were upregulated at PWK 28 in *Cyfip2^+/R87C^* mice compared to WT mice at both PWK 14 and 28, as well as compared to *Cyfip2^+/R87C^* mice at PWK 14. Group 3 lipids were upregulated in *Cyfip2^+/R87C^* mice at both PWK 14 and 28 compared to WT mice at any age. PE, PC, and triacylglycerol (TG) together constituted the major lipid classes in both groups 1 (60%) and 3 (47%). However, diacylglycerol (DG) and cholesteryl ester (CE) were uniquely enriched, accounting for 14% in group 1 and 11% in group 3, respectively. Notably, TG was predominant in group 2, representing 52% of the total lipids. Group 4 included lipid classes that were downregulated in *Cyfip2^+/R87C^* mice at both PWK 14 and 28 compared to WT mice at any age. PE and PC together accounted for 67% of the lipids in group 4, while TG was notably low, comprising only 0.51%. Group 5 represented lipid classes that were upregulated in *Cyfip2^+/R87C^* mice compared to age-matched WT mice. Group 6 consisted of lipid classes upregulated at PWK 28 compared to PWK 14 in WT mice but remained unchanged in *Cyfip2^+/R87C^* mice. This group was notably enriched in hexosylceramide (HexCer). Taken together, these results highlight significant alterations in the proteome, particularly involving mRNA splicing and processing, mitochondrial metabolic pathways, and in the lipidome during seizure progression in *Cyfip2^+/R87C^* mice.

## Discussion

In this study, we employed a comprehensive approach, integrating transcriptomic, ultrastructural, proteomic, and lipidomic analyses for longitudinal deep phenotyping of *Cyfip2^+/R87C^* mice, a model of West syndrome. Notably, these mice exhibit seizure evolution, progressing to spontaneous recurrent seizures in adulthood, which is likely linked to their increased lethality. This underscores the direct relevance of our findings to patients carrying the *CYFIP2* p.Arg87Cys mutation. Given the limited information on the long-term prognosis of these patients, our model suggests that they may also experience seizure evolution, emphasizing the need for long-term follow-up even after the resolution of infantile spasms. Moreover, we identified a complex mechanism involving multiple cell types that exhibit temporally dynamic changes, which may influence one another throughout seizure evolution. Finally, our findings suggest disruptions in brain lipid and metabolic homeostasis as important contributors to seizure evolution, highlighting a potential novel therapeutic target (**Fig 9**). These insights not only enhance our understanding of seizure evolution in West syndrome but also lay the groundwork for exploring shared mechanisms across other epilepsy, neurodevelopmental, and neurological disorders, as discussed further below.

**Fig 9.**
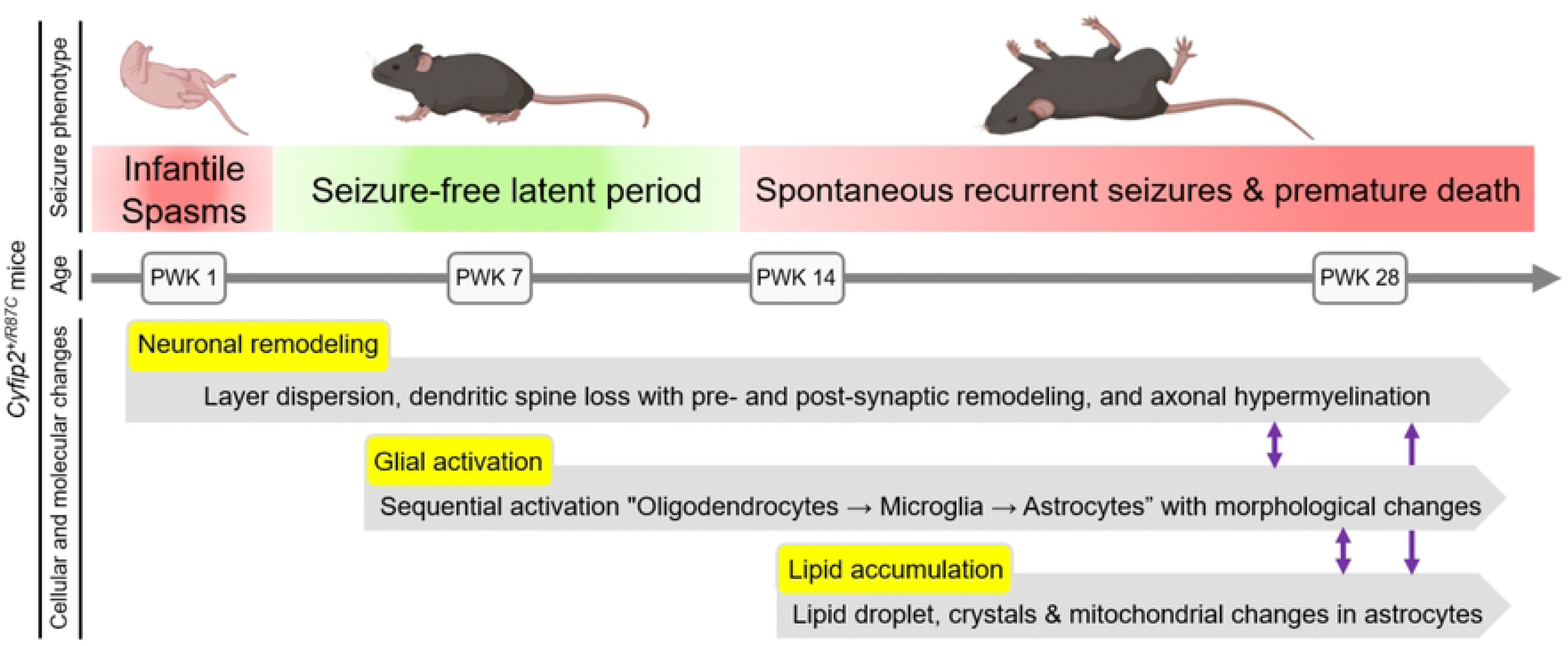
Summary of the seizure evolution in *Cyfip2^+/R87C^* mice. Seizure phenotypes in *Cyfip2^+/R87C^* mice begin with infantile spasms during the neonatal stage (PWK 1), followed by a seizure-free latent period during the juvenile and young adult stages. Around PWK 14, *Cyfip2^+/R87C^* mice start to exhibit spontaneous recurrent seizures, which worsen with age and are associated with premature death. This seizure evolution is accompanied by various temporal cellular and molecular changes in the brain, including alterations in neurons, glial cells, and their lipid and metabolic profiles, which may interact to contribute to the seizure phenotype. The mouse images were obtained from bioRender (https://www.biorender.com/).

The decrease in excitatory synapse number in *Cyfip2^+/R87C^* mice initially seemed counterintuitive, given their spontaneous recurrent seizures, which are typically associated with neuronal hyperactivity. However, detailed molecular and ultrastructural changes in these synapses offer potential explanations. First, dendritic spines, with their narrow neck structure, play a crucial compartmentalizing role in limiting the propagation of biochemical and electrical signals from synapses [38]. When excitatory synapses form directly on the dendritic shaft, bypassing this barrier, it likely becomes easier to activate neurons with each synaptic input, potentially contributing to the observed seizures. Secondly, the dense clustering of hypermyelinated presynaptic inputs may further enhance the activation of postsynaptic neurons. Lastly, although not investigated in detail, the increased expression of presynaptic proteins, such as Synaptotagmin and Synapsin 1, could influence presynaptic vesicle number, distribution, and release probability [39]. Dendritic spine loss observed in epilepsy has often been attributed to damage from excitotoxicity [40], but our results suggest that a more comprehensive and parallel analysis of the molecular and structural properties of the presynapse is needed to better understand the significance of these changes. Further studies, including electrophysiological analysis in *Cyfip2^+/R87C^* mice, are needed to gain a deeper understanding of this type of synaptic remodeling and its role in seizure development.

From the bulk-tissue transcriptomic and proteomic analyses in *Cyfip2^+/R87C^* mice, we observed a downregulation of genes involved in mRNA splicing and processing. Given that CYFIP2 interacts with several RNA-binding proteins associated with these pathways [11, 12], this downregulation may be directly linked to altered CYFIP2 protein function due to the Arg87Cys variant. However, as the downregulation and ASE changes become more evident in *Cyfip2^+/R87C^* mice after PWK 14, when spontaneous recurrent seizures begin, changes in neuronal activity could also contribute to these effects [41]. Notably, *Nrxn2* and *Nrxn3* were identified among the mRNAs showing ASE changes in *Cyfip2^+/R87C^* mice. *Nrxn* genes express various isoforms generated through extensive alternative splicing, which is regulated by neuronal activity and is crucial for synapse formation and specification [35]. Therefore, the ASE changes in *Nrxn2* and *Nrxn3* may contribute to the synapse remodeling in *Cyfip2^+/R87C^* mice, warranting further experimental validation. Furthermore, since ASE changes have been identified in other epilepsy and neurodevelopmental disorders [42], comparing the lists of ASE changes may provide valuable insight into common molecular mechanisms underlying these conditions.

The sequential activation of different glial cell types, beginning with oligodendrocytes, followed by microglia, and then astrocytes, suggests that temporal interactions among these cells play a role during seizure evolution in *Cyfip2^+/R87C^* mice. A recent study has shown that prior activation of microglia is crucial for subsequent astrocytic activation and epileptogenesis in a drug-induced status epilepticus model [43]. Specifically, pharmacological inhibition of microglial activation prevented the development of reactive astrocytes and epileptogenesis in this model. This raises the possibility that sequential activation of glial cell types could be a broader phenomenon during seizure evolution or the latent period of epileptogenesis in various epilepsy types. If so, further investigation is needed to determine whether there is a causal relationship between the activation of different glial cells, including oligodendrocytes, in *Cyfip2^+/R87C^* mice and additional epilepsy models. Understanding these interactions could potentially open multiple therapeutic windows for targeting specific glial cell types to interrupt the progression of seizure evolution and epileptogenesis [44].

Beyond their sequential activations, understanding the roles of different glial cell types in altering neuronal morphology and function, ultimately leading to seizures, is crucial. One clear impact of glial activation on neurons is synapse remodeling or elimination [45], which we directly examined with microglia in the *Cyfip2^+/R87C^* hippocampus.

Oligodendrocytes and astrocytes may also contribute to this process in the *Cyfip2^+/R87C^* mice [46, 47]. Additionally, as mentioned above, the activation of oligodendrocytes in *Cyfip2^+/R87C^* mice likely leads to hypermyelination of axons, which could further influence neuronal activity and connectivity, as previously demonstrated in other epilepsy models [48]. Notably, snRNA-seq-based LR interaction analysis predicted bidirectional crosstalk between CA1-1 and Astrocyte-4 in *Cyfip2^+/R87C^* mice, highlighting their potential influence on target cell gene expression. For instance, CA1-1-induced *Gfap* upregulation in Astrocyte-4 might contribute to the activation of this astrocyte population. Meanwhile, Astrocyte-4-induced regulation of *Adk* and *Grik5* likely affects adenosinergic signaling and glutamatergic synaptic transmission of CA1-1, both of which are strongly associated with epilepsy [49, 50].

We identified changes in LDs in the astrocytes of *Cyfip2^+/R87C^* mice. The various types of LDs observed in these astrocytes may reflect different stages of lipid accumulation. Specifically, LDs enclosed within lysosomes may indicate lipophagy, a form of autophagy that recycles and redistributes lipids in LDs to maintain lipid homeostasis [51]. We suspect that the crystals on or within LDs could be cholesterol crystals, potentially formed due to excess cholesterol storage and LD overload [52]. In either case, these changes suggest abnormal lipid homeostasis in the brains of *Cyfip2^+/R87C^* mice, a hypothesis further supported by single-nucleus transcriptomic data (e.g., upregulation of *Apod* and *Apoe* genes) and lipidomic analyses. Notably, TG, DG, and CE, which are major lipid classes in LDs [53], were upregulated in *Cyfip2^+/R87C^* mice.

In addition to changes in LDs, our proteomic analysis revealed a significant upregulation of mitochondrial proteins involved in energy production and metabolic processes. Morphological alterations in mitochondria were also observed in *Cyfip2^+/R87C^* astrocytes. Given that seizures are highly energy-demanding neuronal activities, it is plausible that the spontaneous recurrent seizures in *Cyfip2^+/R87C^* mice contributed to remodeling mitochondrial properties across multiple cell types in the brain [54]. This remodeling likely disrupted lipid metabolism in astrocytes, including the β-oxidation of fatty acids, ultimately leading to lipid accumulation [55]. Furthermore, hyperactive neurons release excess toxic fatty acids, which are transferred to astrocytic LDs [56]. As a result, spontaneous recurrent seizures may perpetuate a continuous flow of lipids from neurons to astrocytes with impaired mitochondrial function, further disrupting brain lipid homeostasis in *Cyfip2^+/R87C^* mice. Excessive lipid accumulation (lipotoxicity) in astrocytes can further induce mitochondrial dysfunction [57], potentially creating a vicious cycle.

Bidirectionally, mitochondrial metabolic dysfunction and lipid accumulation in astrocytes may exacerbate seizures through various mechanisms, including disruption of the gamma-aminobutyric acid (GABA)-glutamate-glutamine cycle [54, 58]. Notably, changes in astrocytic LDs were more pronounced in PWK 28 compared to PWK 14 *Cyfip2^+/R87C^* mice, suggesting these alterations are more likely associated with seizure severity rather than onset. Moreover, a recent study identified specialized astrocytes, termed lipid-accumulated reactive astrocytes (LARAs), in the brains of temporal lobe epilepsy patients and mouse models [55, 59]. LARAs, potentially due to lipid accumulation-induced gene expression changes, exhibit a reduced capacity to uptake extracellular glutamate, contributing to neuronal hyperactivity. Investigating whether and to what extent astrocytes in *Cyfip2^+/R87C^* mice share functional similarities with LARAs will be an intriguing direction for future research. This could have broad implications for other neurological conditions beyond epilepsy, as mitochondrial metabolic dysfunction and lipid accumulation in astrocytes have recently been recognized as important mechanisms in various neurological disorders through the induction of neuroinflammation and neurodegeneration [60, 61].

In fact, therapeutic approaches targeting the regulation of lipid homeostasis are being explored for various brain disorders, particularly neurodegenerative diseases, at both animal models and clinical levels [62]. For example, treatment with an agonist for liver-X receptor (LXR), a nuclear receptor that regulates cellular lipid homeostasis, not only ameliorated abnormal lipid accumulation in glial cells but also reduced neurodegeneration in a tauopathy mouse model [63]. Our findings, along with the study identifying LARAs [59], suggest that abnormal lipid homeostasis may also play a significant role in the progression of epilepsy. This highlights the potential of targeting lipid homeostasis as a promising therapeutic strategy for epilepsy. The ketogenic diet and its variants, which enhance mitochondrial function and fatty acid oxidation, have demonstrated clinical efficacy in treating medically intractable epilepsy [54]. With a more targeted mechanism, Soticlestat, an inhibitor of the enzyme responsible for converting cholesterol into 24*S*-hydroxycholesterol, had reached phase 3 clinical trials aimed at reducing seizure susceptibility and improving seizure control in Lennox-Gastaut syndrome and Dravet syndrome [64]. Moving forward, further exploration and validation of additional targets for regulating lipid homeostasis will be essential. In this regard, our multi-omic and other comprehensive datasets may provide valuable insights.

We previously demonstrated that Arg87 hotspot variants destabilize CYFIP2 proteins by inducing their ubiquitination and proteasomal degradation [22, 24]. Therefore, given the reduced CYFIP2 protein levels in both lines [15, 22], it was unexpected that *Cyfip2^+/R87C^* mice exhibited much more severe phenotypes than *Cyfip2^+/-^* mice. Transcriptomic analyses of their brains at PWK 1, 7, and 14 revealed distinct age-dependent trajectories, suggesting different underlying mechanisms. While the detailed mechanism of the “toxic gain-of-function” effect of Arg87 variants remains unclear, these findings underscore an important therapeutic concept: selectively targeting and reducing the expression of the *CYFIP2* p.Arg87 mutant allele (i.e., allele-specific) could significantly improve patient outcomes. Approaches like antisense oligonucleotides (ASOs) hold promise for achieving this therapeutic goal [65]. However, once glial activation begins during seizure evolution, direct targeting of the *CYFIP2* mutant allele may be less effective, as CYFIP2 is primarily expressed in neurons rather than glial cells [9, 66]. Therefore, this approach may be most beneficial as an early intervention, possibly around the stage of infantile spasms, before glial activation becomes prominent. Recent studies have explored ASOs as precision therapies for various neurodevelopmental disorders [67]. However, determining the optimal therapeutic window for ASOs and other precision therapies requires a deeper understanding of both the spatiotemporal expression patterns of target genes and the dynamic changes in brain cellular composition throughout disease progression.

In conclusion, our study highlights the value of a longitudinal multi-omic approach in uncovering the complex molecular and cellular dynamics underlying seizure evolution in West syndrome. The resulting datasets represent a valuable resource for future research and may inform broader applications across other epilepsy and neurodevelopmental disorder models.

## Materials and Methods

### Animals

*Cyfip2^+/R87C^* (C57BL/6N background) and *Cyfip2* heterozygous (*Cyfip2^+/-^*, C57BL/6N or C57BL/6J background) mice used in this study have been described previously [15, 22, 68]. Mouse maintenance and related procedures were performed in accordance with the Requirements of Animal Research at Korea University College of Medicine (KOREA-2024-0073) and Korea Advanced Institute of Science and Technology (KA2022-004), as well as with the United States Public Health Service’s Policy on Humane Care and Use of Laboratory Animals. Both male and female mice were used for the experiments.

### Bulk-tissue RNA sequencing and analysis

Bulk-tissue RNA sequencing and analysis were performed as described previously [15, 69, 70]. RNA extraction, library preparation, cluster generation, and sequencing were performed by Macrogen Inc. (Seoul, Korea). RNA samples for sequencing were prepared using a TruSeq Stranded Total RNA LT Sample Prep Kit (Illumina) according to the manufacturer’s instructions. An Illumina’s platform was used for sequencing to generate 101-bp paired-end reads. Transcript abundance was estimated in pseudo-mapping-based mode for the Mus musculus genome (GRCm38) using Salmon (v1.1.0) [71]. Differential gene expression analysis was performed using R/Bioconductor DEseq2 (v1.30.1) [72] by importing the estimated abundance data into R (v.4.1.3) using the tximport package [73]. The *P* values were adjusted for multiple testing with the Benjamini-Hochberg correction. Genes with an adjusted *P* value of less than 0.05 were considered as differentially expressed. Volcano plots were generated using the R ggplot2 (v.3.3.3) package. The Gene Ontology (GO) enrichment analyses were performed using DAVID (https://davidbioinformatics.nih.gov/) software (v2021) [74]. Mouse gene names were converted to human homologs using the Mouse Genome Informatics (MGI) database (http://www.informatics.jax.org/homology.shtml). Gene Set Enrichment Analysis (GSEA) (https://www.gsea-msigdb.org/gsea) [25] was used to determine whether gene expression of a specific set of genes changes in a consistent direction using GSEAPreranked (gsea-4.2.3.jar) module on gene set collections downloaded from Molecular Signature Database (MSigDB) (v7.5.1). GSEAPreranked was applied using the list of all genes expressed, ranked by the fold change and multiplied by the inverse of the *P* value with recommended default settings (1,000 permutations and a classic scoring scheme). The gene sets with a false discovery rate (FDR) of less than 0.05 were considered as significantly enriched. Integration and visualization of the GSEA results were performed using the EnrichmentMap Cytoscape App (version 3.8.1) [75, 76]. The raw data have been submitted to the Gene Expression Omnibus (GEO) repository under the accession numbers GSE292102 (*Cyfip2^+/-^* mice) and GSE292221 (*Cyfip2^+/R87C^* mice).

### Alternative splicing events analysis

We used rMATS v4.1.2 [77] to identify differential alternative splicing events (ASEs) in the cortex and hippocampus of *Cyfip2^+/R87C^* mice compared to WT mice across different postnatal weeks (PWK 1, 7, 14, and 28). To ensure high-confidence results, we filtered out cases where the sum of inclusion junction counts (IJC) and skipping junction counts (SJC) was below 25 in either sample set. Differentially spliced exons were defined using a splicing ratio threshold of |ΔPSI (Percent-Spliced-In)| > 0.2 and a FDR < 0.05.

### RNA purification and real-time quantitative reverse transcription polymerase chain reaction

RNA purification and real-time quantitative reverse transcription polymerase chain reaction (qRT-PCR) were performed as described previously [16, 78]. RNA was purified using the miRNeasy Minikit (QIAGEN, 217004), and cDNA libraries were synthesized using the iScript^TM^ cDNA Synthesis Kit (Bio-Rad, BR170-8891). The target mRNAs were detected and quantified using a real-time PCR instrument (CFX96 Touch, Bio-Rad) with SYBR^TM^ Green Master Mix (Bio-Rad, BT170-8884AP). The results were analyzed using comparative Ct method and normalized against the housekeeping gene *Gapdh*. The primer sequences are listed in S10 Table.

### Western blot analysis

Brain synaptosomal lysate preparation and Western blot analyses were conducted as described previously [79, 80]. Briefly, mouse brains were homogenized in buffered sucrose (0.32 M sucrose, 4 mM HEPES, 1 mM MgCl_2_, 0.5 mM CaCl_2_, pH 7.3) with freshly added protease and phosphatase inhibitors (Sigma-Aldrich, 05892970001 and 04906837001, respectively). The homogenate was centrifuged at 900 g for 10 min, and the resulting supernatant was centrifuged again at 12,000 g for 15 min. The pellet was resuspended in buffered sucrose and centrifuged at 13,000 g for 15 min, resulting in the synaptosome pellet. From each sample, 15–20 μg of protein was loaded onto 4–15% Mini-PROTEAN TGX™ Precast Protein Gels (Bio-Rad, 4561084). The proteins were then transferred to a nitrocellulose membrane (GE Healthcare, 10600001). Tris-buffered saline with Tween 20 (TBST, 0.05%) solution was used for washing, and 5% skim milk in TBST was used for blocking. The Western blot images were captured using the ChemiDoc^TM^ Touch Imaging System (Bio-Rad) and analyzed for quantification using ImageJ software. The primary antibodies used are listed in S11 Table.

### Fluorescence immunohistochemistry

Mice were anesthetized with isoflurane and transcardially perfused with heparinized (20 units/ml) phosphate-buffered saline (PBS), followed by 4% paraformaldehyde (PFA) in PBS. Brains were then extracted and post-fixed overnight in 4% PFA. After post-fixation, brain tissue was washed with PBS and cryoprotected in 30% sucrose in PBS for 48 h. The brain tissues were frozen in O.C.T compound (SAKURA Tissue-Tek, 4583) and coronally sectioned at 60 μm using a cryostat microtome (Leica, CM3050S). Sections were collected from the following approximate Bregma coordinates: mPFC: +2.245 mm to +1.645 mm; hippocampus: −1.555 mm to −2.355 mm. Whole-brain slice imaging was conducted using a Slide Scanner (AxioscanZ1 Fluorescence), and quantification analysis was referenced to the Allen Brain Atlas. For neuronal (NeuN) and glial cell (Olig2, IBA1, and GFAP) markers, we measured intensities instead of performing cell counting, as it was difficult to separately count cells due to their dense clustering and crowding. The mean fluorescence intensity in the mPFC was measured across total layers 1–5 using a 500 µm × 500 µm region of interest (ROI). In the hippocampal CA1 region, the same ROI width was applied to both WT and *Cyfip2^+/R87C^* mice, but the ROI height was adjusted to account for structural differences and to capture dispersed NeuN, particularly in *Cyfip2^+/R87C^* mice. All confocal images were acquired using a ×60 water objective (1.2 numerical aperture). A Z-stack interval of 0.5 µm was used for synapse and BODIPY assessments, while a 0.24 µm Z-stack interval was used for microglia and astrocyte morphology analysis. The primary and secondary antibodies used are listed in S11 Table.

### Imaris quantification

#### Synapse (vGluT1, Homer1)

Synapse numbers were quantified using the Spot tool in Imaris. To assess the number of pre- and post-synaptic puncta in close contact, presynaptic puncta within 0.75 µm of at least one postsynaptic punctum were included in the analysis.

#### Glia cell morphology (IBA1/GFAP)

For IBA1/GFAP morphology, the Filament tool in Imaris was used to trace individual glial cells and quantify branch parameters.

#### Synapse engulfment of microglia (IBA1/Lamp1/Homer1)

The surface of microglia was reconstructed using IBA1 staining (surface detail: 0.2 µm). Lysosomes were reconstructed using the Surface function with Lamp1-positive signals (surface detail: 0.2 µm). The number of Homer1 puncta within Lamp1 vesicles inside microglia was then labeled with the Spot function (spot diameter: 0.5 µm).

#### Lipid droplet (GFAP/Lamp1/BODIPY)

Lipid droplet (LD) expression in microglial lysosomes was analyzed by segmenting the lysosomes using the Surface function to identify Lamp1+/IBA1+ regions. The number of LDs (spot diameter: 1.0 µm) within these regions was then quantified.

### EEG recordings

Electroencephalographic (EEG) recordings and quantifications were performed as described previously [22, 81]. After anesthetizing the mice with a mixture of isoflurane and O_2_ (4%, v/v; 1 NL per min for O_2_ flow and 1.5 L per min for mixed flow) for induction, the mice were positioned on a stereotactic plate (Kopf Instruments, Model 940). The head was fixed, while respiratory anesthesia (2%, v/v) was continuously maintained. Once the skull was exposed, the bregma-lambda axis was aligned with a length of 4.2 mm. Small stainless-steel screws (1 mm × 3 mm) were then implanted on the skull. Two screws were placed in the bilateral prefrontal region (+1.8 mm AP, ±1.0 mm ML from the bregma), and two in the bilateral parietal region (−2.0 mm AP, ±1.8 mm ML from the bregma). Ground and reference screws were positioned on the cerebellum (−1.5 mm AP, ±1.5 mm ML from the lambda). The screws were connected with a connector-housing socket (Hirose Electric, H2021-ND) using soldered-coated stainless-steel wires. The complex was then fixed on the skull using dental acryl. After a week of recovery period, the mice were placed in a white opaque acrylic box (25 × 25 × 35 cm), allowing them to explore freely and to feed *ad libitum* under a 12-h light-dark cycle (13:00-01:00). The EEG recording was performed for 6 days using a Cheetah Data Acquisition System (Neuralynx) synchronized with video recording, 4 days of which were analyzed. The seizure properties were analyzed using Neuraview (Neuralynx), and the representative traces were exported using NeuroExplorer 5 (NeuroExplorer).

### Reflection imaging for visualization of crystals

For the visualization of crystals in *Cyfip2^+/R87C^* mice, tissue sections were fixed and stained with 4’,6-diamidino-2-phenylindole (DAPI), then imaged using Nikon A1R confocal microscope, where a combination of laser reflection and fluorescence confocal microscopy was employed to identify the crystalline materials. Reflectance images were acquired using the confocal system by adjusting the dichroic mirror to the transmitted/reflectance setting (BS20/80), with all light paths set to ‘through’.

### Electron microscopy

Using brain samples from 3 WT and 3 *Cyfip2^+/R87C^* mice at PWK 28, we employed various electron microscopy techniques, generating distinct results for each approach. For sample preparation, the mice were perfused with 2% paraformaldehyde and 2.5% glutaraldehyde in 0.15 M cacodylate buffer with 2 mM Ca^2+^. The brains were post-fixed in the same solution overnight. Fixed brains were sliced into 150 μm-thick sections using a vibratome (Leica, VT1000S) and the hippocampal CA1 region was dissected out.

To assess synaptic density, tissue blocks were prepared following transmission electron microscopy (TEM) protocols as previously described [82]. Briefly, tissue was post-fixed in 2% osmium tetroxide for 1 h, stained en bloc with 1% uranyl acetate, dehydrated, and embedded in Epon 812 resin (EMS). Seventy-nanometer-thick sections were mounted on 200-mesh grids (Gilder Grids, G200N) and post-stained with uranyl acetate followed by lead citrate. For each animal, 20 images were recorded on a Tecnai F20 TEM (Thermo Fisher Scientific) at 3500× magnification and 120kV. A counting frame (4 × 4 μm²) was placed on each image, and synapses with a clear postsynaptic density (PSD) and presynaptic vesicles were counted using stereological methods. Synapses were classified as asymmetric (excitatory) or symmetric (inhibitory) based on their distinct PSD shapes where asymmetric synapses have a thicker PSD and symmetric synapses have a thinner, more uniform PSD. Additionally, the dissected CA1 tissue blocks were prepared for ATUM-SEM imaging using a modified ROTO (Reduced Osmium-Thiocarbohydrazide-Osmium) protocol [15]. This process enabled the three-dimensional visualization of axonal and dendritic structures, as well as the reconstruction of their synaptic connections. The blocks were post-fixed in 2% osmium tetroxide/1.5% potassium ferrocyanide for 1 h, transffered in 1% thiocarbohydrazide for 20 min, and further fixed in 2% osmium tetroxide for 30 min. Samples were incubated in 1% uranyl acetate at 4°C overnight, followed by incubation in lead aspartate solution at 60°C for 30 min. After serial dehydration in ethanol, samples were infiltrated with acetone and embedded in 7% (w/v) conductive Epon 812 resin mixed with Ketjen black powder to enhance electrical conductivity. The specimens, mounted on the aluminum stub, were cured at 60°C for 48 h. After trimming each side of the sample at a 90-degree angle, 50 nm-thick sections were cut with an ultra-Maxi knife (DiATOME, Biel, Switzerland) and collected on plasma-hydrophilized carbon nanotube (CNT)-coated PET tapes (Boeckeler Instruments) using an ATUMtome (Boeckeler Instruments, Inc., Tucson, USA). The sections on CNT tapes were cut into strips, mounted on wafers, and imaged using a Gemini 300 SEM (Carl Zeiss Microscopy GmbH, Oberkochen, Germany). Large-area imaging was conducted using Atlas 5 software (Fibics Incorporated, Ottawa, Canada). Serial SEM imaging was performed at 5 kV with BSD detection, a dwell time of 7 µs, and a resolution of 5 nm × 5 nm per pixel in the X and Y directions, with the images stitched into a single plane. Serial images were aligned using TrackEM2, and one EM stack per brain sample was obtained for both the WT and *Cyfip2^+/R87C^* groups, covering approximately 100 μm × 150 μm in the XY plane with 200 sections at 50 nm each. We reconstructed the dendrites of CA1 pyramidal neurons in the stratum radiatum, followed by reconstruction of the axons forming synapses with these dendrites using an open-source reconstruction software (https://synapseweb.clm.utexas.edu/software-0). Additionally, the surface area of the PSD was measured.

### Correlative light and electron microscopy (CLEM)

To simultaneously detect fluorescence signals of lipid droplets and crystals in EM images, we combined confocal imaging with ATUM-SEM, utilizing the CLEM method. Hippocampal slices, 150 μm thick, were washed three times for 10 min each in 0.15 M sodium cacodylate buffer with 2 mM Ca^2+^, and then incubated with BD493/503 (Thermo Fisher Scientific, D3922) in cacodylate buffer for 30 min at room temperature. The crystal fluorescence signal was detected directly using reflection microscopy, without any additional staining. After staining, DAPI was added to visualize nuclei and blood vessels. In the CA1 region of a coronal brain slice, ROIs containing fluorescence signals were imaged using the Nikon A1R confocal microscope, specifically detecting BODIPY signal or reflected fluorescence. The slices containing the imaged ROIs were dissected and laser-cut into small blocks (PALM MicroBeam, Carl Zeiss). The dissected tissues were processed for heavy metal staining, dehydration, and resin embedding using the modified ROTO protocol [15]. The tissues were then serially sectioned at a thickness of 50 nm using an ATUMtome (Boeckeler Instruments, Inc., Tucson, USA). Low-resolution images of the serial sections were initially acquired to confirm their position relative to marked structures, such as blood vessels and nuclei. Once the target structure was identified, high-resolution (5 nm × 5 nm) serial SEM imaging was performed. Finally, the EM images were matched with the fluorescence images, and the fluorescence-positive structures were three-dimensionally reconstructed.

### Single-nucleus RNA sequencing data analysis

The 10× Genomics Chromium platform was employed to prepare snRNA-seq libraries, enabling high-throughput capture and barcoding of single-nucleus transcripts. Initial data processing and quality control were performed using the Seurat R package (version 5.1.0) [83] to exclude low-quality cells and potential contaminants. Normalization was performed using LogNormalize, and 2,000 highly variable genes were selected. Dimensionality reduction was then conducted via PCA, selecting the number of PCs that explained 90% of the variance. Subsequently, neighbors were identified, and cell clustering was performed using Seurat’s FindClusters function. The results were visualized with UMAP.

Marker genes for each cell cluster were identified through pairwise differential expression analysis using MAST [84], and cell types were annotated by comparing cluster-specific markers to known expression profiles from PanglaoDB and the Human Protein Atlas [85, 86]. Downstream analyses included differential expression analysis (Seurat’s FindMarkers function with MAST method, Log2 fold change > 1, adjusted *P* value < 0.05), gene set enrichment analysis (fgsea R package (version 1.27.0)) [87], Over representation analysis (enrichR (version 0.0.31)) [88], differential abundance analysis of cell neighborhoods (Milo R package (version 3.2) [89], DAseq R package (1.0.0)) [90], Transcription factor analysis (SCENIC R package (version 1.3.1)) [32]. Cell-cell communication analysis was performed using NATMI [91], Connectome [92], logFC Mean [93], and SingleCellSignalR [94] methods. The results from these methods were aggregated and ranked [95] using the liana R package (version 0.1.14) [96], enabling comprehensive prioritization of ligand-receptor interactions. The predicted ligand-receptor pairs were ranked by aggregate rank, and their distribution was plotted to identify potential ligands. Target genes were then identified using the nichenetr R package (version 2.2.0) [33]. The raw data has been submitted to the Gene Expression Omnibus (GEO) repository under the accession number GSE293769.

### Sample preparation for proteomic analysis

Hippocampal tissues were isolated from WT and *Cyfip2^+/R87C^* brains at PWK 14 and 28 and homogenized and lysed with the 1× sodium dodecyl sulphate (SDS) buffer (5% SDS, 50 mM triethylamonium bicarbonate (TEAB), pH 8.5). 500 mg of proteins were reduced and alkylated with final 10 mM Tris (2-carboxyethyl) phosphine (TCEP) and 20 mM indole 3-acetic acid (IAA). Tryptic digestion was performed using an S-trap mini digestion kit (ProtiFi, Huntington, NY) following the provided protocol. In this case, 16.7 mg amounts of mass-spectrometry-grade Trypsin Gold (Promega, Madison, WI) were used for each sample (1:30). The final eluted samples were dried in a speed vacuum. The dried peptides were quantified with a Pierce quantitative colorimetric peptide assay kit (Thermo Fisher Scientific, Rockford, IL). 300 mg of trypsin-digested peptides from each sample were labeled using 18-plex TMT reagent (Thermo Fisher Scientific, Rockford, IL) according to the manufacturer’s instructions. Peptides from WT mice at PWK 14 were labeled with the 126, 127N, 127C, 128N tags and peptides from *Cyfip2^+/R87C^* mice at PWK 14 were labeled with the 128C, 129N, 129C, 130N tags. Peptides from WT mice at PWK 28 were labeled with the 130C, 131N, 131C, 132N tags and peptides from *Cyfip2^+/R87C^* mice at PWK 28 were labeled with the 132C, 133N, 133C, 134N tags. For quality control in TMT experiments, reference samples were prepared by pooling of same amount of all samples and labeled with 134C and 135N TMT tags. Each TMT channel was freshly dissolved in anhydrous acetonitrile (ACN) with a ratio of 0.5:20 (w:v, mg:mL). After incubation for 1 h at room temperature, the reaction was quenched by adding 5 mL of 5% hydroxylamine and incubated for 15 min. Then, samples were combined, dried, and desalted with Pierece peptide desalting spin columns (Thermo Fisher Scientific, Rockford, IL). Total peptides were fractionated into 20 fractions by basic reverse phase liquid chromatography and each eluted peptide sample was vacuum dried. For the liquid chromatographytandem mass spectrometry (LC-MS/MS) analysis, fractionated peptides were diluted by mobile phase A (99.9% water with 0.1% FA).

### LC-MS/MS and protein identification for total proteomic analysis

Dissolved samples were analyzed using an Orbitrap Exploris 480 mass spectrometer (Thermo Fisher Scientific) coupled to an UltiMate 3000 RSLCnano system (Thermo Fisher Scientific) equipped with a nano electrospray source. Samples were trapped on 75 mm × 2 cm C18 precolumn (nanoViper, Acclaim PepMap100, Thermo Fisher Scientific) before being separated on an analytical C18 column (75 mm × 50 cm PepMap RSLC, Thermo Fisher Scientific) for 140 min at a flow rate of 250 nL/min. The mobile phases A and B were composed of 0 and 99.9% acetonitrile containing 0.1% formic acid, respectively. The LC gradient started with 5% B for 20 min. It was ramped to 13% of B buffer for 50 min followed by a gradient from 13 to 25% B buffer for 65 min, then 25 to 95% for 5 min, and 95% for 5 min, after which they were equilibrated for 15 min with 5% B buffer for next run. The voltage applied to produce an electrospray was 2000 V. During the chromatographic separation, the Orbitrap Exploris 480 was operated in data-dependent mode, automatically switching between MS1 and MS2. The full scan resolution was 120 000 at m/z 400. The MS2 scans were performed with HCD fragmentation (37.5% collision energy). Previously fragmented ions were excluded for 30 sec within 10 ppm. The electrospray voltage was maintained at 2.0 kV, and the capillary temperature was set to 275°C.

MS/MS spectra were analyzed using the following software analysis protocol with the Uniprot mouse database. The reversed sequences of all proteins were added to the database for calculation of FDR. ProLuCID [97] in Integrated Proteomics Pipeline software; IP2 (Integrated Proteomics Applications, Inc., San Diego, CA, USA) was used to identify the peptides, with a precursor mass tolerance of 20 ppm, and a fragment ion mass tolerance of 200 ppm. The output data files were filtered and sorted to compose the protein list with two and more peptides assignments for protein identification at a false positive rate less than 0.01. TMT reporter ion analysis was established with Census [98] in Integrated Proteomics Pipeline software within mass error of 20 ppm. Statistical analysis was performed with Perseus [99] software (version 1.6.15) The expressions of proteins between samples were compared using Welch’s t-test with *P* value set at < 0.05. All raw MS data files from this study have been deposited to the repository MassIVE with identifier PXD059911.

### Proteomic data enrichment analysis

Proteomic changes in the hippocampus of *Cyfip2^+/R87C^* mice compared to WT mice were classified into four groups based on relative expression levels with mouse age using clustering analysis. For each protein group, Gene ontology (GO) and Kyoto Encyclopedia of Genes and Genomes (KEGG) pathway enrichment analysis was performed using DAVID (https://davidbioinformatics.nih.gov/) software (v2021) [74]. To further capture the relationships between the enriched terms, Metascape (https://metascape.org/gp/) v3.5.20240901) [100] was applied and a subset of enriched terms has been selected and rendered as a network plot, where terms with a similarity > 0.3 are connected by edges. The top 20 clusters consisting of terms with the highest *P* values were displayed, constrained to no more than 15 terms per cluster and no more than 250 terms in total. Protein-protein Interaction enrichment analysis was also performed using Metascape. Only physical interactions in STRING (physical score > 0.132) and BioGrid are used, and the resultant network contains the subset of proteins that form physical interactions with at least one other member in the list. If the network contains between 3 and 500 proteins, the Molecular Complex Detection (MCODE) algorithm has been applied to identify densely connected network components and the three best-scoring terms by *P* value have been retained as the functional description of the corresponding components.

### Lipidomic analysis

Global lipidomic analysis of hippocampus was performed using ultra-performance liquid chromatography (UPLC) coupled with trapped ion mobility (TIMS) time-of-flight (TOF). Weighed hippocampus tissue samples were placed into homogenizer tubes, each containing three 2.8 mm zirconium oxide beads. For solvent extraction, 1.2 mL of 50% methanol/chloroform (1:1, v/v) was added to each tube, based on a tissue weight of 25 mg, ensuring a consistent weight-to-solvent ratio across all samples, after which the tissues were homogenized. After homogenization, the samples were left to stand at 4°C for 10 min, followed by centrifugation at 12,700 rpm for 20 min. The lower lipid phase was aliquoted into 100 μL, and the solvents were removed using nitrogen gas. The dried extracts were reconstituted in 200 μL of 80% isopropanol, vortexed for 1 min. They were then centrifuged at 12,700 rpm for 10 min at 4°C, and 100 μL of the supernatants transferred to vials for lipidomic analysis.

Untargeted lipid profiling was performed using high-performance liquid chromatography (HPLC; 1290 Infinity; Agilent) coupled with trapped ion mobility spectrometry time-of-flight (TIMS-TOF; Bruker). The column oven and auto-sampler temperatures were maintained at 55°C and 4°C, respectively. The samples were eluted and separated using an Acquity UPLC CSH C18 column (2.1 × 100 mm with 1.7-μm particles; Waters). The LC mobile phase comprised, in positive ion mode, 10 mM ammonium formate and 0.1% formic acid in an acetonitrile/water mixture (6:4, v/v) (solvent A) and 0.1% formic acid in an acetonitrile/isopropanol mixture (1:9, v/v) (solvent B). In negative ion mode, the LC mobile phase comprised 10 mM ammonium acetate in an acetonitrile/water mixture (4:6, v/v) (solvent A) and an acetonitrile/isopropanol mixture (1:9, v/v) (solvent B). Gradient elution began with 40% B, increased to 43% B by 2 min, and then gradually increased to 50% B by 2.1 min. The composition was further increased to 54% B by 12 min, followed by a continuous increase to 99% B by 18 min. The composition was held at 99% B for 0.1 min, it was returned to initial conditions after 18.1 min, and was finally maintained for 2 min until equilibration. The flow rates were set to 400 μL/min. For lipid analysis, 2 μL of lipid extract was injected for positive ion mode, and 4 μL was injected for negative ion mode into the HPLC/TIMS-TOF system. All samples were pooled in equal amounts to generated a quality control (QC) sample. For MS analysis, the mass range was set to m/z 50–1500. The operating parameters were as follows: capillary voltage of 4500 V, end plate offset of 500 V, nebulizer pressure of 2.2 bar for positive ion mode and 2.0 bar for negative ion mode, drying gas flow rate of 10.0 L/min, and source temperature of 220°C.

The acquired data were processed using MetaboScape 5.0 (Bruker) to alignment and peak picking, and identified utilizing the MSDIAL-TandemMassSpectralAtlas-VS69 library. Principal component analysis (PCA) was performed to visualize scatter score plots using SIMCA-P+ software, version 16.0 (Umetrics, Umea, Sweden). Statistical analyses were conducted using Statistical Package for the Social Sciences software, version 28.0 (SPSS Inc., Armonk, NY), and data were plotted using GraphPad Prism, version 8.0 (GraphPad Software, Inc., San Diego, CA). The z-scores were calculated based on the entire sample set. The Kruskal-Wallis U test was conducted to evaluate overall differences of z-scores among the groups. For pairwise comparisons, the Mann-Whitney test was performed, and Bonferroni correction was applied to adjust for multiple comparisons. The results of the lipidomic analysis were deposited in the Korea BioData Station (K-BDS) (BioProject: KAP241486).

## Author Contributions

Youngae.J., J.Y.K., Eunjoon.K., K.J.L., and K.H. contributed to the conception and design of the study. R.M., M.K., G.H.K., Hyojin.K., S.L., Y.Y., H.J.L., S.C., Seungsoo.K., Seoyeong.K., Yukyung.J., Hyewon.K., Y.Z., U.S.K., H.R.K., Y.K., Y.L., W.C., Eunha.K., M.J., G-S.H., J.C., Youngae.J., J.Y.K., Eunjoon.K., K.J.L., and K.H. contributed to data acquisition and analysis. Youngae.J., J.Y.K., Eunjoon.K., K.J.L., and K.H. wrote the paper and prepared the figures.

## Data Availability

All data needed to evaluate the conclusions in the paper are presented in the paper and/or the supporting information.

## Abbreviations

ACN: anhydrous acetonitrile
Adam10: a disintegrin and metalloproteinase domain 10
Adk: adenosine kinase
AMPA: α-amino-3-hydroxy-5-methyl-4-isoxazolepropionic acid
Angpt1: angiopoietin 1
Apod: apolipoprotein D
Apoe: apolipoprotein E
App: amyloid-beta precursor protein
ASE: alternative splicing event
ASO: antisense oligonucleotide
A.U.: arbitrary units
Bdnf: brain derived neurotrophic factor
CE: cholesteryl ester
CLEM: correlative light and electron microscopy
CNT: carbon nanotube
CYFIP2: cytoplasmic FMR1-interacting protein 2
DAPI: 4’,6-diamidino-2-phenylindole
DAVID: Database for Annotation, Visualization, and Integrated Discovery
DEE65: developmental and epileptic encephalopathies 65
DEG: differentially expressed gene
DEP: differentially expressed protein
DG: diacylglycerol
ECM: extracellular matrix
EEG: electroencephalographic
FDR: false discovery rate
GABA: gamma-aminobutyric acid
GEO: Gene Expression Omnibus
GFAP: glial fibrillary acidic protein
GO: Gene Ontology
Grik5: glutamate ionotropic receptor kainate type subunit 5
GSEA: Gene Set Enrichment Analysis
HexCer: hexosylceramide
Homer1: Homer protein homolog 1
HPLC: high-performance liquid chromatography
IAA: indole 3-acetic acid
IBA1: ionized calcium-binding adapter molecule 1
IJC: inclusion junction count
Inhba: inhibin subunit beta A
K-BDS: Korea BioData Station
KEGG: Kyoto Encyclopedia of Genes and Genomes
LARA: lipid-accumulated reactive astrocyte
LC-MS/MS: liquid chromatographytandem mass spectrometry
LD: lipid droplet
LR: ligand-receptor
LXR: liver-X receptor
MBP: myelin basic protein
MCODE: Molecular Complex Detection
MGI: Mouse Genome Informatics
Mif: macrophage migration inhibitory factor
mPFC: medial prefrontal cortex
MSigDB: Molecular Signature Database
NASH: nonalcoholic steatohepatitis
NeuN: neuronal nuclei
Nrxn: Neurexin
Olig2: oligodendrocyte transcription factor 2
OPC: oligodendrocyte precursor cell
PBS: phosphate-buffered saline
PC: phosphatidylcholine
PCA: principal component analysis
Pcdh7: protocadherin 7
PD: postnatal day
PE: phosphatidylethanolamine
PFA: paraformaldehyde
Plpp3: phospholipid phosphatase 3
PPI: protein-protein interaction
Prnp: prion protein
Psap: prosaposin
PSD: postsynaptic density
PSI: Percent-Spliced-In
Pstk: phosphoseryl-tRNA kinase
Ptgds: prostaglandin D2 synthase
PWK: postnatal week
QC: quality control
ROI: region of interest
SCENIC: Single-Cell rEgulatory Network Inference and Clustering
SDS: sodium dodecyl sulphate
Shank3: SH3 and multiple ankyrin repeat domains 3
SJC: skipping junction count
Smox: spermine oxidase
SUDEP: sudden unexpected death in epilepsy
TBST: Tris-buffered saline with Tween 20
TCEP: Tris (2-carboxyethyl) phosphine
TEAB: triethylamonium bicarbonate
TEM: transmission electron microscopy
TG: triacylglycerol
TIMS: trapped ion mobility
TOF: time-of-flight
UMAP: Uniform Manifold Approximation and Projection
UPLC: ultra-performance liquid chromatography
U2af1: U2 small nuclear RNA auxiliary factor 1
vGluT1: vesicular glutamate transporter 1
WAVE: Wiskott–Aldrich syndrome protein family verprolin-homologous protein
WT: wild-type

## Supporting information

**S1 Fig. Representative whole-body images of mice at postnatal week 28 (PWK 28), along with comparisons of body weights.** Scale bar, 2 cm (n = 9 to 36 mice per genotype, unpaired two-tailed Student’s t-test). N = C57BL/6N background, J = C57BL/6J background. \*\*\**P* < 0.001. Data are represented as mean ± standard error of the mean.

**S2 Fig. Imaging and quantification of excitatory synapses in the hippocampus.** A schematic diagram illustrating the hippocampal CA1 region, where the confocal images were acquired. The images on the bottom show examples of converting the hippocampal confocal fluorescence images into spot images using Imaris rendering for WT and *Cyfip2^+/R87C^* mice. Scale bar, 10 μm.

**S3 Fig. Changes in excitatory synapses in the mPFC of *Cyfip2^+/R87C^* mice.** (A) Schematic diagram illustrating the prelimbic medial prefrontal cortex (mPFC), where the confocal images were acquired. The images on the right show examples of converting the mPFC confocal fluorescence images into spot images using Imaris rendering for WT and *Cyfip2^+/R87C^* mice. Scale bar, 10 μm. (B) Representative fluorescence immunohistochemistry images and quantification show changes in excitatory presynaptic (vGluT1) and postsynaptic (Homer1) markers in the prelimbic mPFC of *Cyfip2^+/R87C^* mice compared to WT mice at PWK 14 and 28 (n = 7 to 9 mice per genotype, two-way ANOVA with Šídák’s multiple comparisons test). Images for automated foci counting, obtained using Imaris software, are also shown. A.U. = arbitrary units. \**P* < 0.05. Data are represented as mean ± standard error of the mean.

**S4 Fig. Changes in axonal myelination in the hippocampus of *Cyfip2^+/R87C^* mice.** (A) Representative 3D electron microscopy images of axonal segments in the hippocampal CA1 region from WT and *Cyfip2^+/R87C^* mice at PWK 28. The graph displays the g-ratio (axon diameter/total myelinated fiber diameter) for WT and *Cyfip2^+/R87C^* mice (n = 36 to 41 axons per genotype, unpaired two-tailed Student’s t-test). (B) Representative fluorescence immunohistochemistry images and corresponding quantification show increased myelin basic protein (MBP) mean intensity in the hippocampus of *Cyfip2^+/R87C^* mice compared to WT mice at PWK 28 (n = 8 to 9 mice per genotype, unpaired two-tailed Student’s t-test). DG = dentate gyrus, SO = stratum oriens, SP = stratum pyramidale, SR = stratum radiatum. \*\**P* < 0.01; \*\*\**P* < 0.001. Data are represented as mean ± standard error of the mean.

**S5 Fig. Neuronal and glial changes in the mPFC of *Cyfip2^+/R87C^* mice during seizure evolution.** (A) Representative fluorescence immunohistochemistry images and quantifications show age-dependent changes in the intensities of neuronal and glial cell markers in the prelimbic medial prefrontal cortex (mPFC) of *Cyfip2^+/R87C^* mice compared to WT mice (n = 7 to 9 mice per genotype, two-way ANOVA with Šídák’s multiple comparisons test). The results of the same analysis for *Cyfip2^+/-^* mice (on either the C57BL/6N (N) or C57BL/6J (J) backgrounds) at PWK 28 are also presented (n = 6 to 9 mice per genotype, two-way ANOVA with Šídák’s multiple comparisons test). (B) Changes in the intensities of neuronal and glial cell markers in the hippocampal CA1 region of *Cyfip2^+/-^* mice at PWK 28 (n = 7 to 9 mice per genotype, two-way ANOVA with Šídák’s multiple comparisons test). SO = stratum oriens, SP = stratum pyramidale, SR = stratum radiatum. \**P* < 0.05; \*\**P* < 0.01; \*\*\**P* < 0.001. Data are represented as mean ± standard error of the mean.

**S6 Fig. Changes in morphology of microglia in the mPFC of *Cyfip2^+/R87C^* mice.** Representative 3D images of microglia in the prelimbic mPFC of WT and *Cyfip2^+/R87C^* mice at PWK 14 and 28, processed using Imaris software. Graphs show quantifications from Sholl analysis as well as measurements of branch length, area, points, and diameter (n = 7 to 8 mice per genotype, two-way ANOVA with Šídák’s multiple comparisons test). \**P* < 0.05; \*\**P* < 0.01; \*\*\**P* < 0.001. Data are represented as mean ± standard error of the mean.

**S7 Fig. Changes in morphology of astrocytes in the mPFC of *Cyfip2^+/R87C^* mice.** Representative 3D images of astrocytes in the mPFC prelimbic region of WT and *Cyfip2^+/R87C^* mice at PWK 14 and 28, processed using Imaris software. Graphs show quantifications from Sholl analysis as well as measurements of branch length, area, points, and diameter (n = 7 to 8 mice per genotype, two-way ANOVA with Šídák’s multiple comparisons test). \**P* < 0.05; \*\*\**P* < 0.001. Data are represented as mean ± standard error of the mean.

**S8 Fig. Lysosomal lipid droplets in astrocytes of WT and *Cyfip2^+/R87C^* hippocampus at PWK 14.** Representative 3D images show astrocytes (GFAP), lysosome (Lamp1) within the astrocytes, and lipid droplets (LDs, BODIPY) within astrocytic lysosome (indicated by white arrows in merged images) in the hippocampal CA1 region of WT and *Cyfip2^+/R87C^* mice at PWK 14, analyzed using Imaris software. The Graph shows quantification of the number of LDs within astrocytic lysosomes in WT and *Cyfip2^+/R87C^* mice (n = 7 to 8 mice per genotype, unpaired two-tailed Student’s t-test).

**S9 Fig. Various types of lipid droplets observed in the hippocampal astrocytes of *Cyfip2^+/R87C^* mice.** Serial electron microscopic images of various types of lipid droplets (within lysosome, with or without crystals) in the hippocampal astrocytes of *Cyfip2^+/R87C^* mice at PWK 28. Schematic diagrams for each type of lipid droplet are provided on the left side.

**S10 Fig. Single-nucleus transcriptomic analysis on the hippocampus of *Cyfip2^+/R87C^* mice.** (A) Violin plots showing quality control metrics for each sample: the total number of Unique Molecular Identifier (UMI) counts (left), the number of observed genes (middle), and percent of mitochondrial genes (right). KI, knock-in. (B) UMAP visualization of all hippocampal cells, grouped by sample. Each sample is represented by a distinct color. (C) Volcano plot showing global differential expression (DE) analysis results for all hippocampal cells based on genotype. (D) Subclustering results of neurons. The left panel presents a UMAP visualization of neuronal subtypes, while the right panel shows a dot plot of cell-specific marker gene expression, with dot size representing the percentage of cells expressing the gene and color intensity indicating average expression levels. (E) Subclustering results of excitatory neurons. The left panel displays a UMAP visualization of excitatory neuronal subtypes, while the right panel features a dot plot highlighting cell-specific marker genes. (F) Subclustering results of pyramidal neurons. The left panel presents a UMAP visualization of neuronal subtypes, while the right panel shows a dot plot of cell-specific marker gene expression. (G) Bar plot showing significant GSEA results for CA1-1 compared to CA1-0. The Y-axis represents Gene Ontology Biological Process (GOBP) terms, while the X-axis represents Normalized Enrichment Scores (NES). Color intensity reflects the significance level, represented by −Log10 (adjusted *P* value).

**S11 Fig. Single-nucleus transcriptomic analysis on the glial cells in the hippocampus of *Cyfip2^+/R87C^* mice.** (A) Subclustering results of oligodendrocytes. The left panel presents a UMAP visualization of oligodendrocyte subtypes, while the right panel features a dot plot highlighting the top DEGs for each cluster. (B) Subclustering results of microglia. The left panel presents a UMAP visualization of microglia subtypes, while the right panel features a dot plot highlighting the top DEGs for each cluster.

**S12 Fig. Ligand-target gene predictions between CA1-1 and Astrocyte-4.** (A) Regulatory potential of ligands (Adam10, App, Bdnf, and Inhba) from CA1-1 on twelve predicted target genes expressed in Astrocyte-4. (B) Expression level comparisons of the twelve predicted target genes between Astrocyte-1 and Astrocyte-4. (C) Regulatory potential of Angpt1, App, Mif, and Psap from Astrocyte-4 on their 28 target genes expressed in CA1-1. (D) Over-Representation Analysis (ORA) result for the 28 target genes. (E) Expression level comparisons of the 28 target genes between CA1-0 and CA1-1.

**S13 Fig. Networks of enrichment terms for proteomic analysis of the hippocampus in *Cyfip2^+/R87C^* mice.** (A) Metascape (https://metascape.org/gp/index.html#/main/step1) network of enrichment terms for group B proteins, with nodes sharing the same cluster ID typically positioned close to one another. (B) Metascape network of enrichment terms for group D proteins.

**S14 Fig. PPI networks for group B proteins, highlighting their association with mRNA splicing, processing, and transport.**

**S15 Fig. Heatmap representing lipid species clustered into six groups based on age-dependent relative expression levels in *Cyfip2^+/R87C^* mice compared to WT mice.** Z-scores are significantly different among different groups (n = 9 to 14 mice per genotype, Kruskal-Wallis test).

**S1 Table. List of DEGs for bulk-tissue RNA-seq of *Cyfip2^+/R87C^* mice.**

**S2 Table. List of DEGs for bulk-tissue RNA-seq of *Cyfip2^+/-^* mice.**

**S3 Table. List of DEGs for bulk-tissue RNA-seq of *Cyfip2^+/R87C^* mice at PWK 28.**

**S4 Table.** List of DEGs for snRNA-seq clusters of *Cyfip2^+/R87C^* hippocampus at P28 WK.

**S5 Table.** List of global DEGs for snRNA-seq of *Cyfip2^+/R87C^* hippocampus at P28 WK.

**S6 Table. List of DEPs for proteomics of *Cyfip2^+/R87C^* hippocampus at PWK 14 and 28.**

**S7 Table. List of DEPs clustered into four groups for proteomics of *Cyfip2^+/R87C^* hippocampus.**

**S8 Table. List of alternative splicing event changes in the cortex and hippocampus of *Cyfip2^+/R87C^* mice.**

**S9 Table. List of lipid species clustered into six groups for lipidomics of *Cyfip2^+/R87C^* hippocampus.**

**S10 Table. qRT-PCR primer sequence.**

**S11 Table. Primary and secondary antibody information.**

**S1 Video. Spontaneous seizure captured during video recording in *Cyfip2^+/R87C^* mice at PWK 14.**

**S2 Video. Spontaneous seizure captured during video recording in *Cyfip2^+/R87C^* mice at PWK 28.**

**S3 Video. Tonic-clonic seizure captured during video-EEG recording in *Cyfip2^+/R87C^* mice at PWK 20.**

**S4 Video. Myoclonic jerk captured during video-EEG recording in *Cyfip2^+/R87C^* mice at PWK 20.**

